# Dual role of striatal astrocytes in behavioral flexibility and metabolism in the context of obesity

**DOI:** 10.1101/2023.03.21.533596

**Authors:** Enrica Montalban, Daniela Herrera Moro Chao, Anthony Ansoult, Cuong Pham, Andrea Contini, Julien Castel, Rim Hassouna, Marene Hardonk, Anna Petitbon, Ewout Foppen, Giuseppe Gangarossa, Pierre Trifilieff, Dongdong Li, Serge Luquet, Claire Martin

**Author notes:** corresponding authors (C.M), (S.H.L.), (E.M.). Senior authors (Dongdong Li, Serge Luquet, Claire Martin).

## Abstract

Brain circuits involved in metabolic control and reward-associated behaviors are potent drivers of feeding behavior and are both dramatically altered in obesity, a multifactorial disease resulting from genetic and environmental factors. In both mice and human, exposure to calorie-dense food has been associated with increased astrocyte reactivity and pro-inflammatory response in the brain. Although our understanding of how astrocytes regulate brain circuits has recently flourish, whether and how striatal astrocytes contribute in regulating food-related behaviors and whole-body metabolism is still unknown. In this study, we show that exposure to enriched food leads to profound changes in neuronal activity and synchrony. Chemogenetic manipulation of astrocytes activity in the dorsal striatum was sufficient to restore the cognitive defect in flexible behaviors induced by obesity, while manipulation of astrocyte in the nucleus accumbens led to acute change in whole-body substrate utilization and energy expenditure. Altogether, this work reveals a yet unappreciated role for striatal astrocyte as a direct operator of reward-driven behavior and metabolic control.

## Introduction

Obesity is a major public health problem, which increases the relative risk of a set of pathological conditions (e.g. heart disease, hypertension, type 2 diabetes, steatosis and some form of cancers) (Must et al., 1999; GBD 2015 Obesity Collaborators et al., 2017). Although both genetic and lifestyle factors thoroughly participate in the development of obesity, the contribution of each factor widely varies from individual to individual. Over consumption of highly rewarding high fat, high sugar diet (HFHS) is definitively an identified culprit. While homeostatic circuits located in the hypothalamic-brainstem axis are potent contributors of feeding behaviors, the rewarding nature of food is another powerful drive of feeding (Berthoud et al., 2017). The rewarding aspect of food involves the release of dopamine (DA) within the cortico-mesolimbic system (Berridge, 1996; Alcaro et al., 2007; Björklund and Dunnett, 2007). Consumption of HFHS enhances DA release within the Nucleus accumbens (NAc) and the dorsal striatum (DS) (Lenoir et al., 2007), which in turn influences the striato-hypothalamic circuits promoting food intake (Kenny, 2011; Kempadoo et al., 2013; O’Connor et al., 2015). Alterations in the DA transmission have been shown to be implicated in addictive/compulsive-like ingestive behaviors as well as altered cognitive flexibility (Yang et al., 2018) and reward processing (Koob and Volkow, 2010), two well-established endophenotypes of overweight individuals, which largely depend on striatal processing. It is therefore suggested that, by hijacking the reward system, exposure to palatable hypercaloric diets can switch feeding from a goal-directed and flexible behavior, to an impulsive (Babbs et al., 2013; Adams et al., 2015), inflexible, and ultimately compulsive-like behavior [see (Wang et al., 2001; Johnson and Kenny, 2010; Kenny, 2011; Michaelides et al., 2012)]. In line with this, increasing evidence support that the development of obesity and obesity-related disorders not only results from metabolic dysregulation, but also from dysfunctions of the fronto-striatal circuit, a main substrate for inhibitory behaviors and cognitive control, which can be altered in response to food and associated cues (Stice et al., 2008; Seabrook et al., 2023). However, the cellular and molecular events that underlie the mal adaptive response of the reward system to obesogenic environment remain elusive.

Increasing evidence point to an alteration of astrocytes, the most abundant type of glial cells (García-Cáceres et al., 2019), as a pathophysiological feature of obesity. Astrocytes reactivity, reflected by both morphological and functional remodeling has already been described in the hypothalamus, in response to days or weeks of HFHS exposure, well before fat accumulation and systemic inflammation (Thaler et al., 2012; Clyburn and Browning, 2019). Consumption of enriched diet and the excess of adipose tissue further favor inflammatory cascades associated with secretion of pro-inflammatory signals (Thaler et al., 2012), triggering vascular hyper permeability and maladaptation in both microglia and astrocytes (García-Cáceres et al., 2019).

Despites the physiological evidence that astrocyte are primary target of caloric dense food, it is yet unclear if they play a dominant role in the cognitive and metabolic defect associated with obesity. In the current study, we show that long-term exposure to HFHS leads to profound changes in striatal astrocytes states and activity, associated with loss of synchrony in neuronal activity and impairs mice reversal learning. Second, we show that selective manipulation of striatal astrocyte through chemogenetic approaches helps reinstate neural network coordination. Third, we identified a neuroanatomical distinction by which activation of astrocytes in the dorsal striatum can directly rescue HFHS diet-induced cognitive dysfunction while manipulating astrocytes activity in the Nucleus accumbens exert a dominant control onto whole-body substrate utilization and energy expenditure.

## Results

### High-fat diet-induced obesity leads to reactive astrocytes in both the Nucleus Accumbens and the Dorsal Striatum

Previous studies have demonstrated that HFHS exposure results in reactive astrocytes (Douglass et al., 2017) and alters astrocytic calcium signals in the hypothalamus (Herrera Moro Chao et al., 2022). Anatomical and functional studies have suggested a functional heterogeneity within the striatum, with the ventral striatal regions more likely to be involved in goal directed behaviors, and the dorsal subdivisions rather related to motor control and habits development (Kravitz and Kreitzer, 2012; Lee et al., 2012). Therefore, we explored the distinctive astrocytic adaptations in both the DS and the NAc. Mice were exposed to HFHS for a minimum of 3 months (**Fig-1A**), leading to a significant increase in fat mass compared to chow fed littermates (**Fig-1A**). In both NAc and DS, exposure to HFHS diet enhanced immunoreactivity of the structural protein glial fibrillary acidic protein (GFAP) (**Fig-1B,C,F**), a proxy of increased astrocyte reactivity (Escartin et al., 2021). In HFHS fed mice, the increase in GFAP signal intensity was also accompanied by a decrease in the sphericity of the segmented GFAP positive regions in both the DS and the NAc (**Fig-1D,G**), indicating an effect of HFHS diet on astrocytes morphology. In HFHS-fed groups, the total surface of GFAP staining was significantly increased in the NAc indicating an increase of astrocytic coverage, (**Fig-1E**) while unchanged in the DS (**Fig-1H**). Altogether, these data indicate a functional heterogeneity in the striatal astrocyte response to HFHS diet exposure.

**Figure-1.**
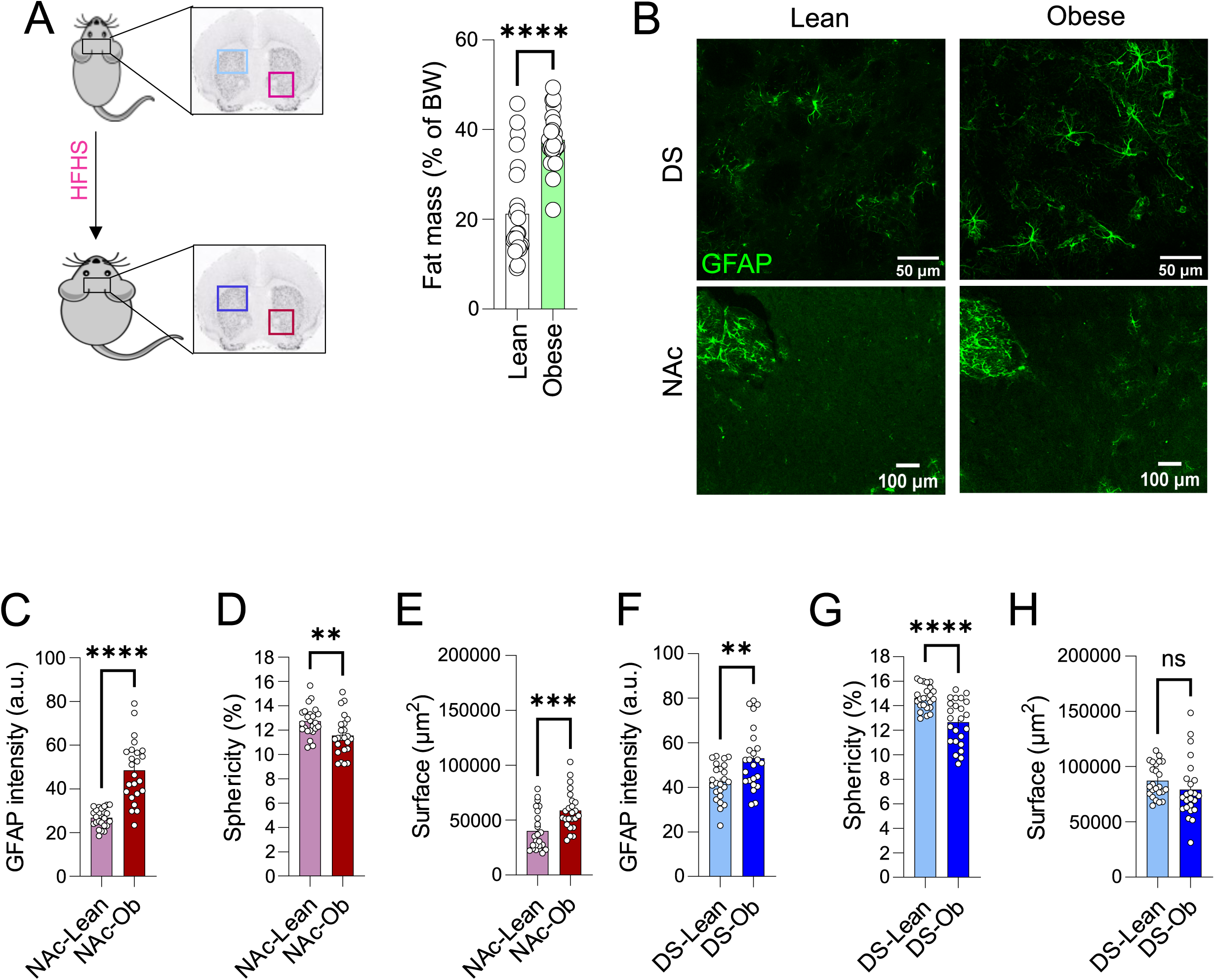
DIO increases glial-fibrillary acidic protein (GFAP) immunoreactivity in the DS and the NAc. **A Left-**Schematic representation of the protocol. **Right**-After 190 days of HFHS diet obese mice showed a significant increase in fat mass as compared to lean. Unpaired Mann-Whitney test ****p<0.0001, n =24 **B** Confocal images representative of GFAP immunoreactivity in the DS and the NAc of lean and DIO mice. **C-H** In the NAc and the DS, DIO increases relative expression of GFAP immunoreactivity compared to lean (**C, NAc; F, DS**). DIO also results in a decrease of astrocytes sphericity (**D-NAc**, unpaired t-test p=0.0064/**G-DS** unpaired t-test p<0.0001). Total surface of astrocytic coverage was decreased by DIO in NAc, **E-NAc**, unpaired t-test p<0.0001, while was left unchanged in the DS **H-DS** unpaired t-test p=0.2496). All data are expressed as mean ± SEM. n = 24, 6 mice in each group. **I-L** DIO increases Ca2+ strength and decreases the overall temporal correlation of astrocyte Ca2+ signal intensity in Glast-GCaMP6 mice expressing GCaMP selectively under the GLAST promoter (For NAc, Lean: n = 643 active regions, 24 slices, 6 mice; Obese: n = 586 active regions, 19 slices, 4 mice; for DS, in Lean, n = 204 active regions, 11 slices, 4 mice; Obese, n = 178 active regions, 8 slices, 3 mice).

### Chemogenetic manipulation of DS astrocytes affects spiny projections neurons activity

The finding that HFHS exposure triggers structural and functional changes in striatal astrocytes led us to assess metabolic and behavioral consequences of astrocytic manipulation in the striatum of lean and obese mice. We first probed the consequence of chemogenetic (Designed Receptors Exclusively activated by Designer Drug)-mediated manipulation of DS astrocytes on DA signaling mediated by pharmacological intervention onto dopamine 1 receptor (D1R) and dopamine 2 receptor (D2R). Lean and obese mice expressing the CRE recombinase under the control of the astrocytes-specific promoter Aldehyde dehydrogenase family 1, member L1 (Aldh1l1-Cre) (Cahoy et al., 2008) were stereotactically injected with Cre-dependent pAAV-EF1α-DIO-hM3Dq-mCherry in the DS allowing for the astrocyte-specific expression of the Gq-coupled receptor (DS^hM3Dq^) (**Supp. Fig-1**). Astrocytic-specific targeting was confirmed by co-immunolocalization of the mCherry signal in striatal astrocytes with the astrocyte’s marker GFAP (**Supp. Fig-1A**). Next, to validate the DREADD-induced Ca^2+^ signaling in astrocytes, we co-expressed the Ca^2+^ indicator GCaMP6f using Cre-dependent AAV vector. Intraperitoneal injection (IP) of the DREADD ligand Clozapine N-Oxide (CNO, 0.6 mg/kg) led to significant increase of astrocytic Ca^2+^ activity as assessed *in vivo* through fiber photometry recording of DS GCaMP6-based fluorescence (**Supp. Fig-1B, C**). As a functional readout, we observed that Gq-DREADD-mediated manipulation of astrocytes in the in DS astrocyte did not alter hyper locomotion triggered by a single injection of the D1R agonist (SKF-81297) (**Supp Fig-1D**), while the cataleptic effects induced by the D2R antagonist haloperidol (0.5 mg/kg) was significantly decreased in response to the DREADD ligand CNO. Interestingly, this effect was further enhanced in obese mice (**Supp. Fig-1E-F**).

### Diet-induced obesity leads to increased temporal correlation of astrocyte Ca^2+^ signals in the DS

Next, we investigated how HFHS exposure impacts onto spontaneous astrocytic Ca^2+^ dynamics, an important feature of astrocyte signalling (Agulhon et al., 2008; Khakh and McCarthy, 2015). To do so, we used mice expressing the genetically encoded Ca^2+^ sensor GCaMP6f under the astrocyte-specific promoter of the glutamate-aspartate transporter (*Slc1a3*, GLAST) (Glast-GCaMP6f), (Pham et al., 2020; Herrera Moro Chao et al., 2022). Wide-field imaging of striatal astrocytes in acute brain slices of lean and obese Glast-GCaMP6f mice showed that exposure to HFHS does not affect Ca^2+^ spontaneous activity in the DS (**Fig-2A**). However, in the DS, obesity associates with an increased temporal correlation of astrocyte Ca^2+^ signals, as showed by calculating the paired Pearson’s coefficient, a correlation coefficient between individual Ca^2+^ signals reflecting signal synchronicity. While the distribution of temporal correlation of Ca^2+^ signals from all active domains appeared bimodal in lean mice, suggesting that astrocyte Ca^2+^ signals are segregated into two populations of asynchronous temporal features, this distribution is right shifted in obese mice indicating increased temporal correlation (**Fig-2B**). These results show that beyond a change of astrocyte Ca^2+^ signaling intensity, exposure to HFHS changes the temporal organization of astrocyte activation. Because astrocytes are tightly linked with synaptic activity, it is likely that this shift also affects neuronal synchronization.

**Figure-2.**
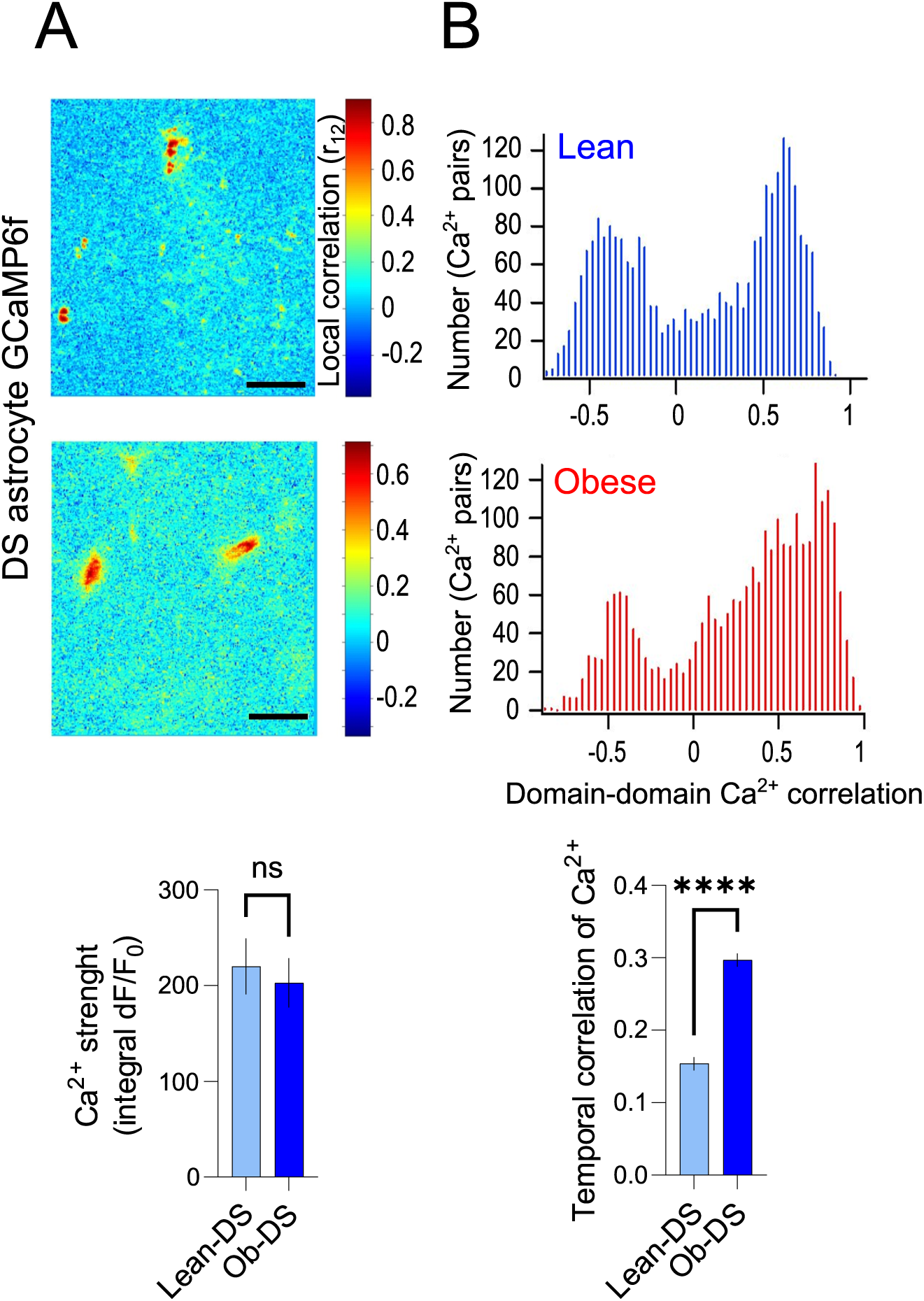
DIO alter region specific astrocytic Ca^2+^ activity in the DS. DIO decreases the overall temporal correlation of astrocyte Ca^2+^ signal intensity in Glast-GCaMP6 mice expressing GCaMP selectively under the GLAST promoter in the DS. Lean, n = 204 active regions, 11 slices, 4 mice; Obese, n = 178 active regions, 8 slices, 3 mice. **A**. Representative pseudo-images of the GCaMP6 fluorescence projection of the spontaneous Ca^2+^ activity in the NAc of lean (top) and obese (middle) mice. Histogram data (bottom) are expressed as mean +/− SEM. Scale bar: 20 µm **B**. Distribution of temporal correlations of Ca2+ responses of all paired active domains (as an estimation of global synchronization) in lean (top) and obese (middle). Overall Ca^2+^ strength data (bottom) are expressed as mean +/− SEM.

### DS astrocytes chemogenetic manipulation in obese mice rescues neuronal synchronization

Using brain slice preparation for GCaMP6f monitoring we observed that obesity had little impact on the overall strength of neuronal Ca^2+^ signals (**Supp. Fig-2A**), but significantly decreased temporal correlation pattern of neuronal events and reduced on average their correlation level as compared to lean animals (**Fig-3B**), suggesting that obesity compromised the synchrony in the DS neuronal network. To further explore the impact of obesity on astrocyte-neuron communication in the DS, we examined the effect of astrocytes chemogenetic manipulation on neuronal Ca^2+^ signals *ex vivo*. C57Bl6 mice received a mixture of viral vectors allowing for simultaneous expression of GCaMP6f in neurons (AAV-synapsin-GCaMP6f) and hM3Gq in astrocytes (AAV-GFAP-hM3Gq-mCherry) (**Fig-3A**). Next, we analyzed neuronal time courses during Gq-DREADD astrocyte activation in lean and obese animals. We first validated that coincident activation of neuronal populations can enhance their synchrony. To do so, we used glutamate whose receptors are abundantly expressed in DS neurons (Montalban et al., 2022) and applied a concentration (30µM) that targets perisynatpic mGluR and/or NMDA receptors, hence mimicking the activation of glutamate receptors targeted by Gq-DREADD astrocyte activation. Application of glutamate did synchronize the Ca^2+^ events in GCaMP6f-expressing DS neurons, as reflected by simultaneous fluorescence rises (**Supp. Fig-2B, C**) and the right shifted distribution of the temporal correlations (**Supp. Fig-2D, E**).

**Figure-3.**
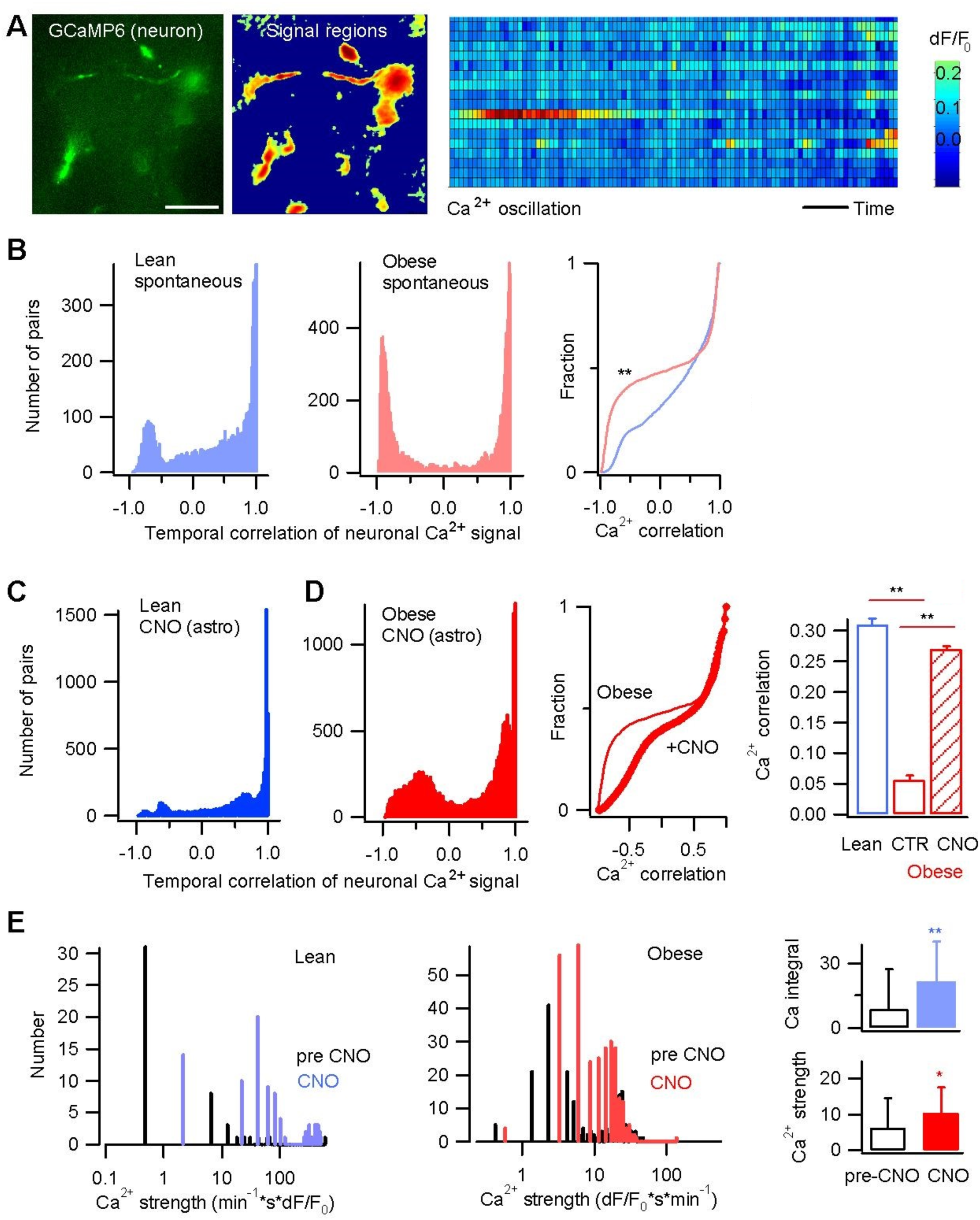
Activation of astrocytes augments neuronal activity synchrony in the DS in obese mice. **A.** Neuronal spontaneous activity was recorded by Ca^2+^ imaging with GCaMP6. *Left*, temporal projection of GCaMP6 fluorescence, scale bar, 20 µm; *middle*, identified regions displaying Ca^2+^ oscillations; *right*, raster plot showing GCaMP6 fluorescence fluctuations over time indicating the spontaneous Ca^2+^ signals. Scale bar, 50 µm. Temporal bar, 10 s. **B.** Distribution of temporal correlation of neuronal Ca^2+^ signal between lean and obese mice. Temporal correlation was derived from the Pearson’s correlation coefficients calculated between all pairs of individual Ca^2+^ signals. 6131 signal pairs for four mice for lean, and 11796 pairs from three mice for obese condition. Wilcoxon rank sum (Mann-Whitney) test, p = 1.65 × 10^−6^, h = 1, stats = [zval: 18.58, ranksum: 6.1067 × 10^7^]. **C.** Distribution of the temporal correlation of neuronal Ca^2+^ signals in response to astrocyte Gq DREADD activation in lean mice (9199 signal pairs from three mice). Lean and obese as referenced from **B**. **D.** In obese mice, astrocyte Gq DREADD activation by CNO enhanced the temporal correlation (synchrony) of neuronal Ca^2+^ signals (20326 signal pairs from three mice; p = 6.97 × 10^−125^, h = 1, stats = [zval: −23.7691, ranksum: 1.7042 × 10^8^]). **E.** Activation of astrocyte Gq DREADD enhances neuronal Ca^2+^ intensity in lean (p = 1.06 × 10-24, h = 1, stats = [zval: −10.2608, ranksum: 4438]) and in obese (ranksum, p = 0.032, h = 1, stats = [zval: −2.1403, ranksum: 82174]) mice. The Ca^2+^ strength derived from normalized temporal integral was compared for control (pre CNO) and CNO application phase (Lean, 89 responsive regions, 3 slices, 3 mice; Obese: 294 regions, 5 slices, 3 mice).

### Activation of DS astrocytes rescues neuronal synchronization defect associated with obesity

We then examined whether Gq-mediated activation of DS astrocytes could modulate neuronal activity profile as assessed by GCaMP6f activity. Bath application of CNO significantly increased the level of temporal correlation between neuronal Ca^2+^ signals, leading to a right shift of the correlation distribution (**Fig-3C-D**). Notably, the impairment of neuronal Ca^2+^ signals synchrony associated with obesity was largely restored by Gq-DREADD-mediated astrocytes activation (**Fig-3D**), along with overall enhancement of neuronal activity (**Fig-3E**). To further confirm this effect, Aldh1l1-Cre mice were co-injected with Gq-DREADDs or mCherry control viruses (DS^mCherry^ and DS^hM3Dq^) and AAV-synapsin-GCaMP6f to target neurons. As previously observed, the strength (**Supp. Fig-3A-B**) and temporal correlation (**Supp. Fig-3C**) of DS neuronal Ca^2+^ signals were increased by both glutamate bath application and CNO-mediated DREADD manipulation of astrocytes. Together, these results show that the signal synchronization of DS neurons is dampened in obese mice, but can be restored by selective activation of striatal astrocytes.

### Obesity-associated impairment in cognitive flexibility can be rescued by selective activation of striatal astrocyte

We next explored the functional outcome of DS astrocytes manipulation onto obesity-induced cognitive alteration. Reversal learning is a form of cognitive flexibility highly dependent on to the integrity of the DS and that was shown to be impaired in human and rodent obesity(Foldi et al., 2021; Montalban et al., 2023). Neuroimaging studies in humans show that reversal learning requires the integrity of the ventral prefrontal cortex and the DS (Jocham et al., 2009). Previous studies already showed that activation of astrocytes in the DS facilitate the switch from habitual to goal directed behavior in lean mice in a operant conditioning paradigm (Kang et al., 2020). We first evaluated if DS-dependent flexible behavior was altered in obese mice. To do so, lean and obese mice of matched age were tested in a food-cued T-maze, in which mice learnt to locate the baited arm with no external cues, using an egocentric strategy (Oliveira et al., 1997; Watson and Stanton, 2009; Baudonnat et al., 2013) followed by a reversal learning task, in which locations of the baited and non-reinforced arms are inverted (**Fig-4A**). While no differences were observed during the learning phase, obese mice displayed impaired ability to relearn the new location of the baited arm during reversal task (**Fig-4A**). Their performances did not reach criterion even after 3 sessions of reversal test (**Supp. Fig-4**) whereas lean mice reached 80% of correct choice during the first reversal session (**Fig-4A**).

**Figure-4.**
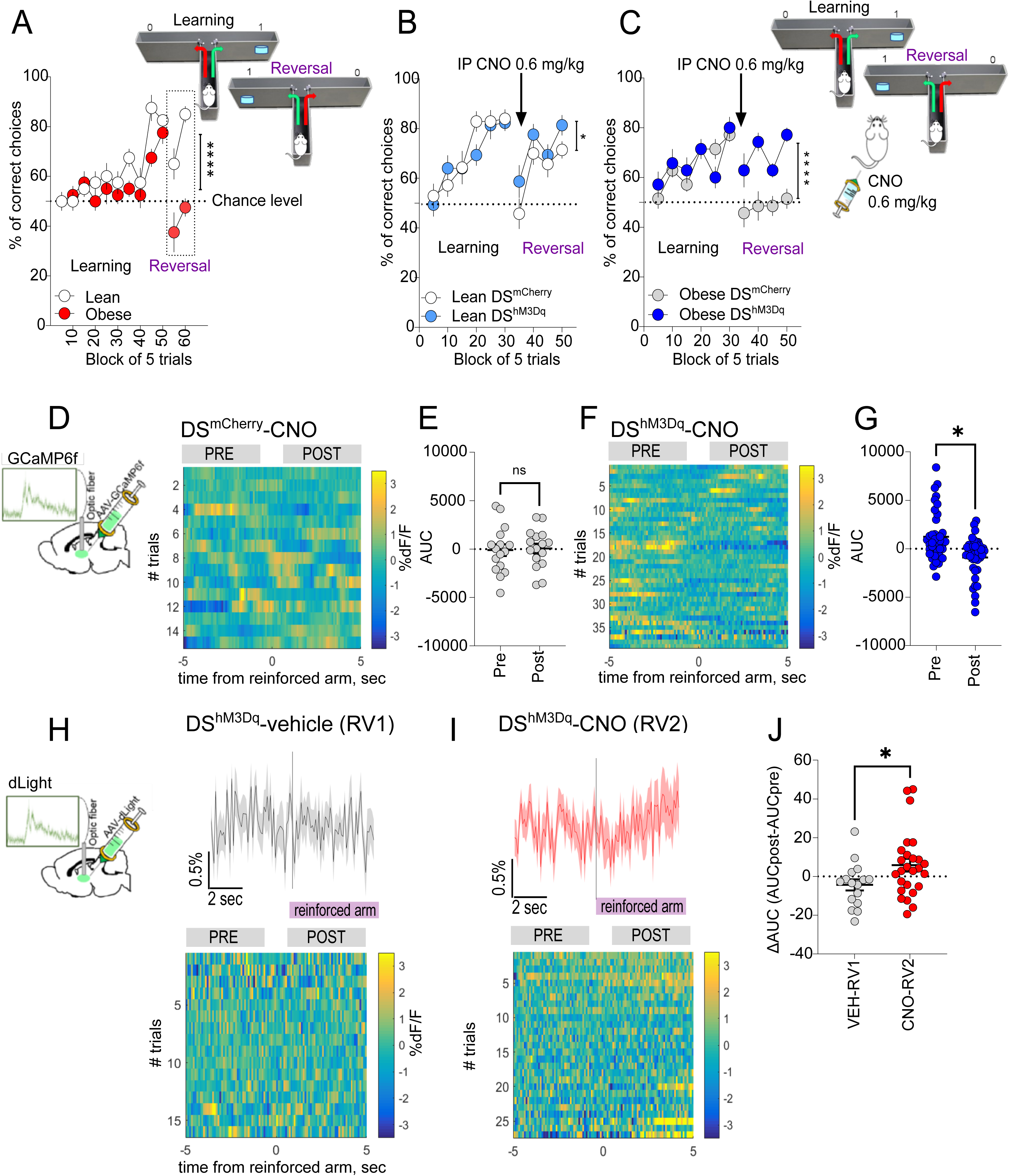
Effect of DS astrocytes activation on the reversal learning in a T-maze paradigm in Aldh1l1-cre lean and obese mice. A) Right behavioral paradigm. Left Performances of Aldh1l1^DS-mCherry^ lean and obese mice were compared for learning and reversal learning skills. A significant between group difference in the reversal phase indicate a decreased flexibility in obese as compare to lean mice. Reversal phase, two-way ANOVA Column Factor F (1, 14) = 33.80 P<0,0001 Data are expressed as mean ± SEM. n= 8. **B.** DS^mCherry^ and DS^hM3Dq^ lean mice were trained in a T-maze and injected with CNO before the reversal phase. CNO injections slightly increases flexibility in DS^hM3Dq^ mice as compared to control. Group Factor F (1, 27) = 5,125 P=0,0318, Data are expressed as mean ± SEM. n= 14-15. **C** Astrocytes activation before the reversal phase in obese DS^hM3Dq^ mice restore the behavioral performances. Two-way ANOVA: Group Factor (1, 12) = 35,54 P<0,0001 Data are expressed as mean ± SEM. n= 7. **D-G** Neuronal Ca^2+^ activity was evaluated during reversal learning by fiber photometry in the DS of DS^mCherry^ and DS^hM3Dq^ mice co-injected with a virus expressing GCaMP6f in DS neurons. Each mouse was injected with CNO 30 minutes before the test and recorded during the T-maze session. **D,F** Peri-event heat map of the single trials of DS^mCherry^ and DS^hM3Dq^ mice respectively, aligned to the time when mice attained the baited arm. **E,G** Plot of area under the curve (AUC) during the baited arm exploration vs before turning in the baited arm (4 s each, indicated by horizontal grey bars) in mice treated with CNO (n = 16 and 39 trials for DS^mCherry^ and DS^hM3Dq^ mice respectively). Statistical analysis two-tailed Mann-Whitney test, p = 0.74 for DS^mCherry^ and p=0.0126 for DS^hM3Dq^ mice. **H-J** DA transmission was evaluated by fiber photometry during reversal learning in DS of DS^mCherry^ and DS^hM3Dq^ mice co-injected with a virus expressing dLight-1 in DS neurons. Each mouse was recorded twice with an interval ≥ 1 day (Reversal day 1 and Reversal day 2), 30 min after receiving either vehicle (**Veh**) or misoprostol (**CNO**, 0.06 mg.kg^−1^, i.p.). **H,I** Peri-event heat map of single trials of mice injected with Veh or CNO aligned to the time when mice attained the baited arm. Plot of area under the curve (AUC) during the baited arm exploration minus the AUC before turning in the baited arm (4 s, horizontal bars) in mice treated with Veh (RV1) vs CNO (RV2), Statistical analysis two-tailed Mann-Whitney test, p = 0.0360 (n = 16 and 29 trials for Veh and CNO respectively).

Next, we assessed the consequence of Gq-DREADDs activation of DS astrocyte on reversal learning in lean and obese Aldh1l1 DS^mCherry^ and DS^hM3Dq^ mice. Reversal learning was assessed in response to CNO injection 30 minutes before the first trial of the reversal phase (**Fig-4B, C**). Importantly, while activation of Gq-DREADD in DS astrocytes in lean mice led to a small though significant increased performance (**Fig-4B**), CNO injection in obese DS^hM3Dq^ led to an almost complete restoration of reversal learning during the reversal phase (**Fig-4C**). Our results indicate that DS astrocytes activation during reversal learning was sufficient to restore obesity-induced impairment in cognitive flexibility.

### Astrocyte-mediated restoration of flexible behavior in obese mice is associated with changes in both neuronal activity and dopamine transmission *in vivo*

In order to link the behavioral output with bulk neuronal activity in the DS upon Gq-DREADD astrocytes activation *in vivo*, we recorded neuronal activity using Ca^2+^ sensor coupled with fiber photometry during reversal learning. Our analysis showed that astrocytes activation during reversal learning was accompanied by a small decrease of neuronal activity when the animal enters the baited arm in obese DS^hM3Dq^ as compared to DS^mCherry^ (**Fig-4D-G**).

Several studies indicate that obesity and HFHS exposure enhances DA signaling in both humans (Volkow and Wise, 2005) and rodents (Johnson and Kenny, 2010; Tellez et al., 2013), and recent studies point to a role of astrocytes in regulating the level of DA release in the striatum (Roberts et al., 2022). Since DA transmission regulates behavioral flexibility (Izquierdo et al., 2017), we investigated the role of DS astrocytes in DA transmission during the reversal learning. Obese DS^mCherry^ and DS^hM3Dq^ mice were co-injected with a viral vector bearing the genetically-encoded DA sensor dLight1 in the DS (AAV-CAG-dLight1.1)(Patriarchi et al., 2018). Fiber photometry recording of dLight1-mediated signal was used as a proxy of DA dynamics in the DS. Mice were first recorded during a reversal learning after being injected with vehicle (RV1, **Fig-4H-J**) and, next, during a second reversal learning after being injected with CNO (RV2, **Fig-4H-J**). Our analysis showed that astrocytes activation during reversal learning potentiated DA transmission when the animal entered the baited arm (**Fig-4I, J**).

Overall our data showed that in DS astrocytes activation in obese mice restores reversal learning impairments in relation with i) an overall decrease in neuronal activity and ii) an increase in DA transmission when entering the new-bated arm during reversal learning.

### Diet induced obesity leads to increased strength and decreased temporal correlation of astrocyte Ca^2+^ signals

Converging evidence point to a central role of the ventral part of the striatum in the regulation of food intake (Sears et al., 2010; O’Connor et al., 2015; Thoeni et al., 2020), glucose metabolism (Ter Horst et al., 2018) and whole body substrate utilization (Montalban et al., 2023). Hence, we next considered a possible role of NAc astrocytes in the physiology and pathophysiology of energy balance in obesity.

First, we investigated the effect of HFHS exposure on spontaneous activity of NAc astrocytes in GLAST-GCaMP6f mice. Interestingly, we found that contrarily to the DS, exposure to HFHS led to a significant increase in NAc astrocytic Ca^2+^ spontaneous activity (**Fig-5A**). Moreover, as for the DS, lean mice showed a bimodal Ca^2+^ distribution suggesting that astrocyte Ca^2+^ signals are segregated into two populations of asynchronous temporal features (**Fig-5B**). However, in contrast to the DS, exposure to HFHS diet led to a left monomodal distribution in the NAc, indicating a decrease in astrocytes synchronization (**Fig-5B**). We next examined the effect of astrocyte activation in Aldh1l1-cre mice that received intra NAc delivery of Cre-dependent viral vectors encoding for Gq DREADD and GCaMP6f. We observed a significant increase in NAc astrocyte Ca^2+^ levels following CNO bath application in lean mice (CNO, 10µM) (**Supp. Fig-5A, B**). This observation led us to hypothesize that Gq-DREADD-mediated increase of astrocytic Ca^2+^ in the NAc of obese mice would have little effect as compared to stimulation of astrocytes in lean mice. We used pharmacology to assess whether astrocytes manipulation would influence behavioral response to agonist and antagonist of D1R and D2R. We observed that NAc astrocytes activation opposed SKF-81297 (3mg/kg) induced hyperlocomotion in lean mice, while this effect was dampened in obese mice (**Supp. Fig-5D, E**). In contrast to DS, astrocytes activation in the NAc did not trigger any significant effects in the cataleptic response to the D2R antagonist haloperidol (0.5 mg/Kg) (**Supp. Fig-5C**) further supporting segregated function of NAc vs DS astrocyte.

**Figure-5.**
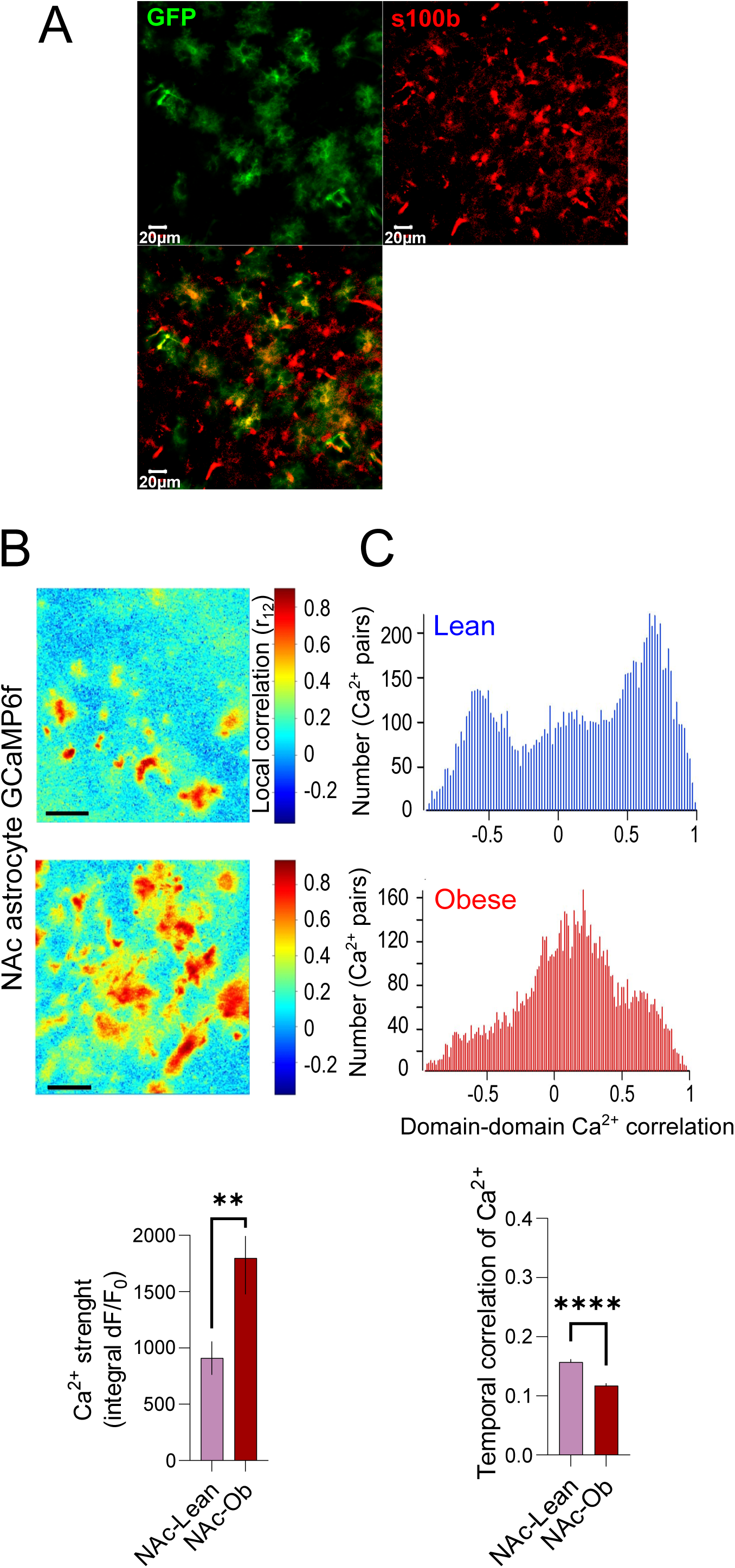
DIO increases Ca2+ strength and decreases the overall temporal correlation of astrocyte Ca^2+^ signal intensity in Glast-GCaMP6 mice expressing GCaMP selectively under the GLAST promoter in the NAc. Lean: n = 643 active regions, 24 slices, 6 mice; Obese: n = 586 active regions, 19 slices, 4 mice. **A** Coronal brain slice showing the colocalization of GFP signal (green) and S100b (red) immunostaining in GCaMP-GLAST mice. **B**. Representative pseudo-images of the GCaMP6 fluorescence projection of the spontaneous Ca^2+^ activity in the NAc of lean (top) and obese (middle) mice. Histogram data (bottom) are expressed as mean +/− SEM. **C**. Distribution of temporal correlations of Ca2+ responses of all paired active domains (as an estimation of global synchronization) in lean (top) and obese (middle). Overall Ca2+ strength data (bottom) are expressed as mean +/− SEM.

### Astrocytes activation in the Nucleus accumbens impacts on peripheral substrate utilization

We then assessed changes in metabolic efficiency in lean and obese Aldh1l1-Cre mice co-injected with Gq-DREADDs or mCherry viruses (NAc^mCherry^ and NAc^hM3Dq^) and AAV-synapsin-GCaMP6f by monitoring indirect calorimetry in response to CNO-mediated astrocytes manipulation (**Fig-6A**). In lean mice, acute stimulation of NAc astrocytes only marginally affected feeding (**Fig-6B**), but promoted a significant decrease in respiratory exchange ratio (RER, VCO_2_/VO_2_) indicative of substrate being used with RER=1 for carbohydrate and RER=0.7 for lipids (**Fig-6C**). Correlation studies indicated that such decrease in RER significantly correlated with food consumption in Aldh1l1 NAc^Gq^ group (**Fig-6D**). In accordance, the calculated whole body fat oxidation (Fat Ox) confirmed that acute activation of astrocytes in the NAc led to a shift towards lipid-based substrate (**Fig-6E**). While whole body metabolism (**Supp. Fig-6**) remained unaffected by activation of DS astrocyte in both lean and obese DS^mCherry^ or DS^hM3Dq^ mice, chemogenetic manipulation of NAc astrocytes also resulted in a decrease of energy expenditure (EE) (**Fig. 6F**). This effect was independent from the mice lean body mass (**Fig-6G**) and locomotor activity, which are not different between groups (**Fig-6H**). In obese mice however, activation of astrocytes in the NAc did not alter either food intake, RER, FatOx or EE (**Fig-6I-L**), further supporting the notion that obesity led to maladaptive response in astrocytic control of metabolism.

**Figure-6:**
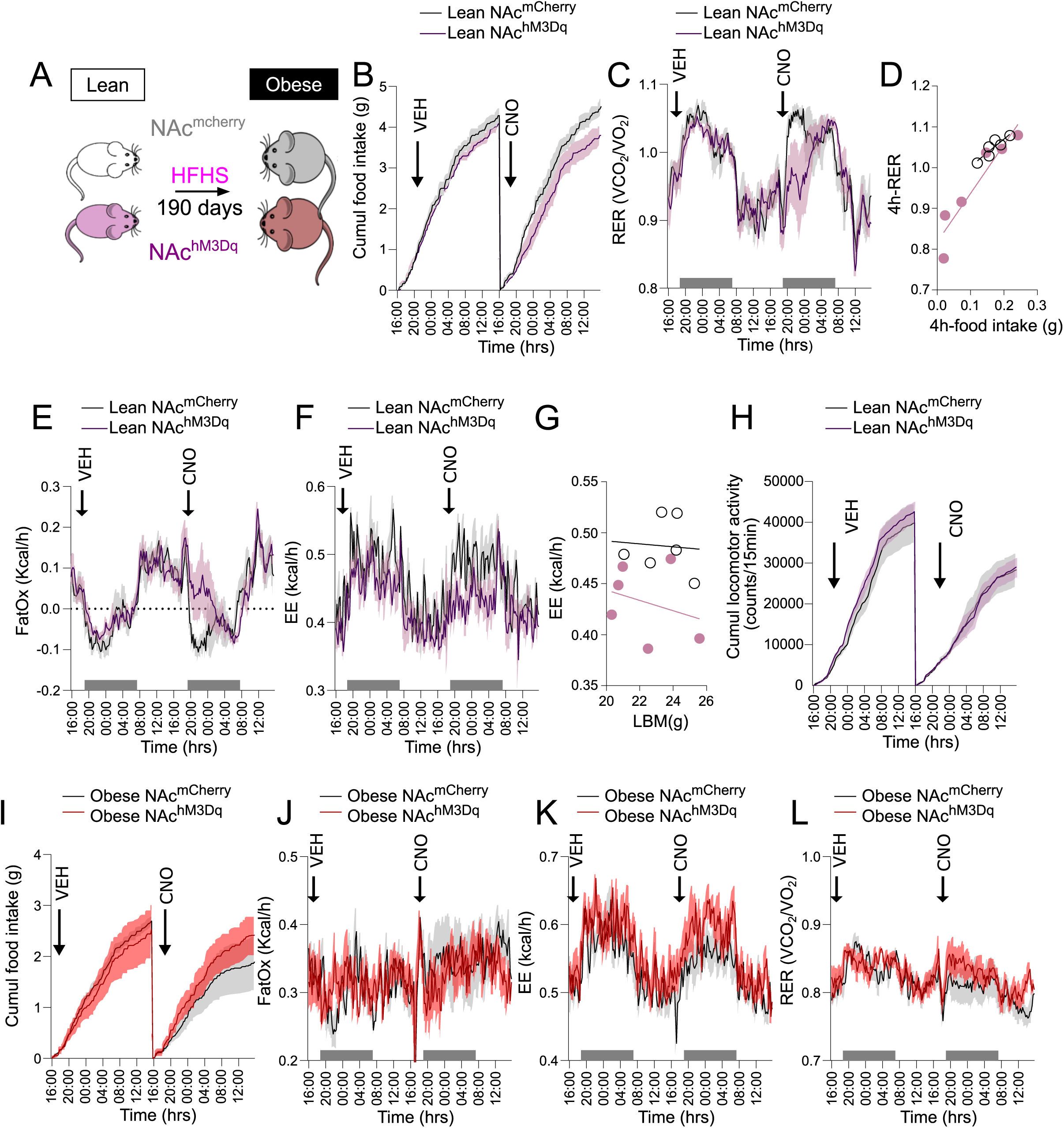
Metabolic consequences of chemogenetic activation of Gq signaling in astrocytes of the NAc in lean and obese mice. **A.** Schematic representation of the DIO paradigm. Astrocytes activation in in NAc^hM3Dq^ mice fed with chow diet does not alter food intake **B** but decreases respiratory exchange ratio (RER) in NAc^hM3Dq^ mice fed with chow diet **C**. RER correlate with caloric intake for both NAc^mCherry^ and NAc^hM3Dq^ mice **D**. Astrocytes activation in lean NAc^hM3Dq^ mice increases fatty acid oxidation **E** and decreases energy expenditure (EE) **F**. EE does not correlate with LBM in neither NAc^mCherry^ mice or NAc^hM3Dq^ mice. **G**. Locomotion is not impacted by astrocytes activation **H**. Metabolic parameters are not impacted by CNO injection in obese NAc^mCherry^ and NAc^hM3Dq^ mice fed with HFHS diet. Cumulative caloric intake, FatOx, energy expenditure (EE) and respiratory exchange ratio (RER) are not significantly modified by CNO injection **I-L**. (N = 6 mice each group; VEH: vehicle).

## Discussion

In the context of the obesity pandemic, the striatum has attracted increasing attention, as energy-rich diets are known to promote reward dysfunctions by altering DA transmission within both NAc and DS. Such alterations can lead to maladaptive habits formation, food craving, inability to cut down food intake and, ultimately to body weight gain. However, while the role of striatal neurons is actively investigated, the contribution of striatal astrocytes in the development of metabolic defects is still largely overlooked. Here, we tested the hypothesis that i) in a diet-induced obesity paradigm striatal astrocytes could be a major target of nutrient overload and that ii) manipulation of astrocytes in DS or NAc could restore behavioral and metabolic alterations induced by obesity. Consistently, we show that obesity induces anatomically-specific change in astrocyte reactivity characterized by substantial alteration in their morphology in both NAc and DS, recalling the modifications observed in the hypothalamus (Thaler et al., 2012). Further, we showed that HFHS consumption translates in an anatomically restricted change in overall Ca^2+^ strength in NAc astrocytes but not in DS astrocytes. In contrast, while temporal correlation in astrocytes Ca^2+^ events was similar in NAc and DS in lean mice, HFHS exposure led to a shift towards a monomodal Ca^2+^ events distribution in the NAc (decreased synchronization), and a significant increase in temporal correlation as compared to lean mice in the DS. Those findings highlight that the functional heterogeneity of astrocytes may reflect different kind of activations among or within brain regions according to their interactions with different subpopulations of neurons (Khakh and Sofroniew, 2015).

Obesity is a condition characterized by both metabolic and behavioral alterations. Among the latter, non-flexible behavior is a symptomatic dimension which is well characterized in obese subjects. Here we measured reversal learning, a dimension known to be particularly affected in obese subjects, relying on an egocentric-based strategy, a process highly dependent on the integrity of DS (van Elzelingen et al., 2022), and that requires the integrity of the ventral prefrontal cortex and DS (Jocham et al., 2009). Using chemogenetics, we showed that activation of DS-astrocytes in lean and obese mice facilitate flexible behavior during the reversal learning phase of a T-maze task. In obese mice, astrocyte activation was sufficient to restore learning flexibility during reversal. These data are in line with a key role of the DS astrocytes in the switch from habitual to goal directed behavior (Kang et al., 2020), and highlight a central role of astrocytes for the long-term consequences of obesity. Using both chemogenetics, GCaMP6f and d-Light based imaging of neural activity and DA transmission *in vivo*, we showed that in obese mice the reinstatement of a flexible behavior under astrocytes activation parallels with a general decrease in neuronal activity in the DS together with an increase in DA transmission during the choice phase, i.e. when mice are entering the rewarded arm. Dysfunctional DA transmission is associated to several psychiatric pathologies characterized by alterations in flexible behavior (Insel et al., 2010; van Elzelingen et al., 2022). These data confirm that reestablishing the DA transmission within the DS correlates with a gain in the ability to adapt its behavior (Leroi et al., 2013) and are in line with previous reports showing that astrocytes are active players in DA signaling in the striatum (Martín et al., 2015; Corkrum et al., 2020)

In line with this, we found that i) obesity is accompanied with a sharp decrease of Ca^2+^ dynamics synchronization in spiny projection neurons SPNs of the DS, that ii) astrocytes activation can restore neural Ca^2+^ event synchronicity, and that iii) in obese mice glutamate application can mimic chemogenetic activation of astrocytes by restoring neural Ca^2+^ events synchrony. These data extend previous works showing that astrocytes can modulate neuronal networks excitability and switch dynamic states *ex vivo* and *in vivo* (Fellin et al., 2004; Poskanzer and Yuste, 2011, 2016; Oliveira and Araque, 2022). Synchronized activity is a defining feature of the nervous system that correlates with brain functions and behavioral states. Several brain diseases are associated with abnormal neural synchronization (Uhlhaas and Singer, 2006). In the striatum, rearrangement in neuronal synchronization plays a key role in habitual learning (Howe et al., 2011; Thorn and Graybiel, 2014; Smith and Graybiel, 2016), hence it is tempting to propose that modulation of synchrony by astrocytes translates into the modifications of behavior that we observed in our experiments. Gq-DREADDs activations and concurrent Ca^2+^ increase can have many consequences on is the so-called the tripartite synapse (Araque et al., 1999). Astrocytes release and uptake neuroactive molecules that could impact both pre- and postsynaptic neuronal functions (Leybaert and Sanderson, 2012; Orellana et al., 2016; Savtchouk and Volterra, 2018)-an effect that have been already suggested in the NAc (D’Ascenzo et al., 2007). Astrocytes are also known to shape synaptic activity and communication by precisely buffering the level of extra synaptic glutamate concentration (Isaacson, 1999; Martin et al., 2012). In this study we observed that in obese but not lean mice, glutamate application increased neuronal synchrony. Since glutamate or CNO application, and CNO glutamate co-application resulted to comparable effects in acute slices from obese mice, a possibility would be that in the DS, obesity would result in a deregulation of glutamate reuptake from astrocytes, an effect that could be rescued by astrocytes activation. Hence, at mechanistic level, our data suggest a central role for astrocyte in controlling neural Ca^2+^ events synchrony and DA transmission. Altered regulation of glutamate in obesity is a mechanism reminiscent of our recent study that depicted a key role for hypothalamic astrocyte in the regulation of neurons firing ability, energy expenditure and glucose metabolism through the control of ambient glutamate (Herrera Moro Chao et al., 2022). We found that obesity was associated with exacerbated astrocyte Ca^2+^ activity and blunted astrocyte-selective excitatory Amino-Acid Transporters (EAATs)-mediated transport of glutamate (Herrera Moro Chao et al., 2022). Since a large portion (∼ 80%) of glutamate released is actively recaptured by astrocyte through Glutamate transporter 1 (GLT-1) and EAATs transporters, it is expected that striatal astrocyte will have a key role in the control of glutamate in the striatum. Indeed, it was recently demonstrated that glutamate transporter in the astrocytes was in control of Hebbian plasticity expression in the SPNs in the DS (Valtcheva and Venance, 2016).

In line with emerging evidence that point at the connection between the DA circuits and metabolic control (Montalban et al., 2023), we found that, activation of NAc astrocytes in lean mice led to significant shift towards lipids substrate utilization. Given the connection between the NAc and hypothalamic nuclei involved in metabolic control, notably the lateral part of the hypothalamus (LHA) (Stratford and Kelley, 1999; Sears et al., 2010; O’Connor et al., 2015; Thoeni et al., 2020) it is formally possible that NAc astrocytes activity indirectly impede onto a subset of neurons projecting to the LHA, with consequences on hypothalamic control of energy expenditure and lipids metabolism (Farzi et al., 2018). This hypothesis is consistent with previously proposed role for a hypothalamic-thalamic-striatal axis in the integration of energy balance and food reward (Kelley et al., 2005). In obese mice, activation of astrocytes of the NAc failed to affect energy metabolism suggesting impaired astrocyte-neural coupling induced by obesity, possibly through astrocyte over activity. Indeed, our *ex vivo* studies showed that Gq-coupled hM3Dq activation in astrocyte leads to increases Ca^2+^ signals similar to the one observed in obese conditions. Therefore, it is tempting to hypothesize that activation of astrocytes in the NAc of lean mice would mimic at least in part some of the obesity metabolic dimensions similarly to what has been observed in hypothalamic astrocyte (Herrera Moro Chao et al., 2022).

Our data support the notion that the physiological outcome arising from astrocyte manipulation will strongly depend on the anatomical localization and the way astrocyte interacts with different subpopulations of neurons (Khakh and Sofroniew, 2015). Indeed, while CNO-mediated activation of Gq-coupled DREADD astrocytes in the DS had marginal effect on metabolic efficiency, chemogenetic activation of the NAc astrocyte in lean mice decreased energy expenditure and sustained change in nutrient portioning interpedently from caloric intake. In the same concept, chemogenetic manipulation of DS astrocytes decreased the cataleptic effects induced by the D2R antagonist haloperidol but did not affect hyperlocomotion response to D1R agonist SKF-81297 suggesting a bias action of astrocyte towards D2R-bearing cells. This result was mirrored in the NAc in which hyperlocomotion response to D1R agonist but not cataleptic response to D2R antagonist was affected by the activation of hM3Dq in the NAc astrocyte. These data directly point at a segregated action of astrocyte in the dichotomic action onto specific neuronal population and DA receptor signaling likely due to the selective and anatomically defined properties of astrocyte-neurons communication. For instance in the DS, two distinct subpopulation of astrocytes have been identified that communicate selectively with D1R or D2R-SPNs (Martín et al., 2015). In addition to this intrinsic diversity in astrocyte-neurons communication, our study highlights that exposure to caloric dense food differently affects astrocyte-neuron communication in NAc and DS. While the consequence of NAc astrocyte activation onto metabolic efficiency observed in lean mice was mitigated by obesity, the cognitive improvement associated with DS astrocyte activation was magnified in obese mice. Here too, the differential impact of high fat feeding might reflect the intrinsic diversity in adaptive response to metabolic signals in DS vs NAc astrocyte, neurons, or both astrocyte-neurons tandem.

In conclusion, this study provides a ground for a more astrocentric vision of diet and obesity induced alteration in cognitive and metabolic function and open new therapeutic avenue in which striatal astrocytes could represent potential target to correct behavioral and metabolic diseases. However, in order to fully harvest the therapeutic potential of an astrocytic-specific target strategy there is a critical need to further expand our knowledge in molecular specificity and mechanism that sustain astrocyte-neuron dialogue in both physiological and pathophysiological condition based on their anatomical distribution.

### Limitations of the study

Due to paucity of tools readily available to characterize DS or NA astrocyte diversity, our study could not provide a more detailed description of the specific features of astrocytes involved in the described mechanism. Further, while changes in astrocytic or neural Ca^2+^ events are indicative of cell response they most likely coexist with other intracellular changes that are not accounted for in our study. Further studies are needed to establish the molecular transmitters, metabolite or metabolic pathways that are engaged in astrocyte-neurons connection and which of them represent the best target to leverage as future strategy to cope for diet-induce metabolic and cognitive disease.

--end--

## AUTHORS CONTRIBUTION

E.M. initiated, developed and supervised the project, designed and performed experiments, analyzed and interpreted the data, prepared figures and wrote the original draft. CM supervised and developed the research, designed and performed in vivo experiments, analyzed and interpreted the data, prepared figures and participated in the writing of the manuscript with the help of the co-authors. SHL provided the initial conception of the project, secured and administered funding, provided guidance for experimental design and data interpretation and contributed to the writing of the manuscript with the help of the co-authors. DL designed and performed ex-vivo calcium imaging experiments, analyzed and interpreted the data, prepared figures and participated in the writing of the manuscript. PT contributed to analysis and interpretation of the data and writing of the manuscript. GG contributed to the design and provided inputs to in vivo experiments and discussed the data. DHMC and CP performed ex-vivo calcium imaging experiments, AC, AP, PT contributed to fiber photometry experiments. DHMC, AA, JC, RH, MH, EF contributed to in vivo experiments and immunohistochemistry.

## ACKNOWLEDGEMENTS

We acknowledge funding supports from l’Agence Nationale de la Recherche (ANR) ANR-19-CE37-0020-02, ANR-20-CE14-0020, and ANR-20-CE14-0025-01 “AstrObesity” and from the Fondation pour la Recherche Médicale (FRM) FRM Project #EQU202003010155. We thank the Université Paris Cité, CNRS, INRAE and Université de Bordeaux. EM was supported by a post-doctoral fellowship from the FRM and awarded by the « Fondation des Treilles ». « La Fondation des Treilles, créée par Anne Gruner-Schlumberger, a pour vocation de nourrir le dialogue entre les sciences et les arts, afin de faire progresser la création et la recherche. Elle accueille des chercheurs et des créateurs dans son domaine des Treilles (Var) www.les-treilles.com ». We thank Olja Kacanski for administrative support, Isabelle Le Parco, Aurélie Djemat, Daniel Quintas, Magguy Boa Ludovic Maingault and Angélique Dauvin for animals’ care and Florianne Michel for genotyping. We acknowledge the technical platform Functional and Physiological Exploration platform (FPE) of the Université Paris Cité, CNRS, Unité de Biologie Fonctionnelle et Adaptative, F-75013 Paris, France, the viral production facility of the UMR INSERM 1089 and the animal core facility “Buffon” of the Université Paris Cité/Institut Jacques Monod. We thank the animal facility of IBPS of Sorbonne Université, Paris.

## METHODS

### Experimental models and subject details

#### Animal studies

All animal protocols were approved by the Animal Care Committee of the University of Paris (APAFIS #2015062611174320), or the Institut Biologie Paris Seine of Sorbonne University (C75-05-24). Twelve to fifteen-week-old male Aldh1-L1-Cre (Tg(Aldh1l1-cre) JD1884Htz, Jackson laboratory, Bar Harbor, USA), male C57BL/6J (Janvier, Le Genest St-Isle, France) or male GCaMP6f/Glast-CreERT2 (Pham et al., 2020) mice were individually housed at constant temperature (23± 2°C) and submitted to a 12/12h light/dark cycle. All mice had access to regular chow diet (Safe, Augy, France) and water ad libitum, unless stated otherwise. Additionally, age matched C57BL/6J, GCaMP6f/Glast-CreERT2 or Aldh1-L1-Cre mice groups were fed with either chow diet or high-fat high-sugar diet (HFHS, cat n. D12451, Research Diets, New Brunswick, USA) for twelve to sixteen weeks. Body weight was measured every week and body weight gain was estimated as the difference of body weight in week one of HFHS diet consumption to twelve to sixteen weeks after HFHS diet exposure.

#### Viral constructs

Designer receptor exclusively activated by designer drugs (DREADD) and GCaMP6f viruses were purchased from http://www.addgene.org/, unless stated otherwise. pAAV-EF1α-DIO-hM3Dq-mCherry (2.4×1012 vg/ml, Addgene plasmid #50460-AAV5; http://www.addgene.org/50460/; RRID: Addgene_50460), pAAV-EF1α-DIO-mCherry (3.6×1012 vg/ml, Addgene plasmid #50462-AAV5; http://www.addgene.org/50462/; RRID: Addgene_50462), pAAV-EF1a-DIO-hM3D(Gq)-mCherry was a gift from Bryan Roth (Addgene plasmid # 50460; http://n2t.net/addgene:50460; RRID: Addgene_50460). pAAV-CAG-Flex.GCaMP6f.WPRE (3.15×1013 vg/ml, working dilution 1:10, Addgene plasmid #100835-AAV5; http://www.addgene.org/100835/; RRID:Addgene_100835) was a gift of Douglas Kim and GENIE Project. pAAV-GfaACC1D.Lck-GCaMP6f.SV40 (1.53×1013 vg/ml, working dilution 1:5, Addgene plasmid #52925-AAV5; http://www.addgene.org/52295/; RRID: Addgene_52925) was a gift of Baljit Khak. pAAV-CAG-dLight1.1 was a gift from Lin Tian (Addgene viral prep # 111067-AAV5; http://n2t.net/addgene:111067; RRID: Addgene_111067)

#### Surgical procedures

For all surgical procedures, mice were first intraperitoneal (ip) injected with the analgesic Buprenorphine (Buprecare, 0.3 mg/kg, Recipharm, Lancashire, UK). 30 minutes after the injection mice were rapidly anesthetized with isoflurane (3%), intraperitoneal (ip) injected with the analgesic Buprenorphine (Buprecare, 0.3 mg/kg, Recipharm, Lancashire, UK) and Ketoprofen (Ketofen, 10 mg/kg, France) and maintained under 1.5% isoflurane anesthesia throughout the surgery.

Stereotaxic surgery. Male Aldh1-L1-Cre+/−, Aldh1-L1-Cre−/− and male C57BL/6J mice were placed on a stereotactic frame (David Kopf Instruments, California, USA) and bilateral viral injections were performed with 0.6ul in DS (stereotaxic coordinates: L = +/−1.75; AP = +0.6; V = −3.5, and −3 in mm), or 0.3ul in NAc (L=+/−1; AP=+1.55, V=-4.5) at a rate of 50 nl.min^−1^. The injection needle was carefully removed after 5 min waiting at the injection site and 2 min waiting half way to the top. Mice recovered for at least 3 weeks after the surgery before being involved in experimental procedures.

#### Behavioral assays

##### Haloperidol-induced catalepsy

Mice were injected with haloperidol (0.5 mg.kg^−1^, i.p.). Catalepsy was measured at several time points, 45-180 min after haloperidol injection. Animals were taken out of their home cage and placed in front of a 4-cm elevated steel bar, with the forelegs upon the bar and hind legs remaining on the ground surface. The time during which animals remained still was measured. A behavioral threshold of 180 seconds was set so the animals remaining in the cataleptic position for this duration were put back in their cage until the next time point.

##### T-maze

Mice were tested for learning and cognitive flexibility in a gray T maze (arm 35-cm length, 25-cm height, 15-cm width). All mice were mildly food deprived (85-90 % of original weight) for 3 days prior to starting the experiment. The first day mice were placed in the maze for 15 min for habituation. Then, mice underwent 3 days of training with one arm reinforced with a highly palatable food pellet (HFHS, cat n. D12451 Research Diet). Each mouse was placed at a start point and allowed to explore the maze. It was then blocked for 20 seconds in the explored arm and then placed again in the starting arm. This process was repeated 10 times per day. At the end of the learning phase all mice showed a > 70 % preference for the reinforced arm. The average number of entries in each arm over 5 trials was plotted. Two days of reversal learning followed the training phase during which the reinforced arm was changed and the mice were subjected to 10 trials per day with the reward in the arm opposite to the previously baited one.

##### SKF-induced locomotor activity

Mice were placed in an automated online measurement system using an infrared beam-based activity monitoring system (Phenomaster, TSE Systems GmbH, Bad Homburg, Germany). After 1 day of habituation, mice were first i.p. injected with CNO (0.6 mg/Kg) and 30 minutes after with SKF-81297 (3 mg/kg), and placed back in the chamber for at least 80 minutes. Locomotion was recorded using an infrared beam-based activity monitoring system Phenomaster, TSE Systems GmbH, Bad Homburg, Germany).

#### Fiber photometry

Aldh1-L1-Cre mice were anaesthetized with isoflurane and received 10 mg.kg-1 intraperitoneal injection (i.p.) of Buprécare® (buprenorphine 0.3 mg) diluted 1/100 in NaCl 9 g.L-1 and 10 mg.kg-1 of Ketofen® (ketoprofen 100 mg) diluted 1/100 in NaCl 9 g.L-1, and placed on a stereotactic frame (Model 940, David Kopf Instruments, California). We unilaterally injected 0.6 µl of virus (pAAV.Syn.Flex.GCaMP6f.WPRE.SV40, Addgene viral prep #100833-AAV9, titer ≥ 1013 genome copy (GC).mL-1, working dilution 1:5) or d-Light1 (pAAV-CAG-dLight1.1, Addgene viral prep # 111067-AAV5, titer ≥ 7×10¹² vg/mL, working dilution 1:1) into the DS (L = +/−1.5; AP = +0.86; V = −3.25, in mm) at a rate of 50 nl.min-1. The injection needle was carefully removed after 5 min waiting at the injection site and 2 min waiting half way to the top. Optical fiber for calcium imaging into the striatum was implanted 100 µm above the viral injection site. A chronically implantable cannula (Doric Lenses, Québec, Canada) composed of a bare optical fiber (400 µm core, 0.48 N.A.) and a fiber ferrule was implanted 100 µm above the location of the viral injection site in the DS (L = +/−1.75; AP = +0.6; V = −3.5, and −3 in mm). The fiber was fixed onto the skull using dental cement (Super-Bond C&B, Sun Medical). Real time fluorescence emitted from the calcium sensor GCaMP6f expressed by astrocytes with the Aldh1-L1-Cre receptor was recorded using fiber photometry as described in (Berland et al., 2020). Fluorescence was collected in the DS using a single optical fiber for both delivery of excitation light streams and collection of emitted fluorescence. The fiber photometry setup used 2 light emitting LEDs: 405 nm LED sinusoidally modulated at 330 Hz and a 465 nm LED sinusoidally modulated at 533 Hz (Doric Lenses) merged in a FMC4 MiniCube (Doric Lenses) that combines the 2 wavelengths excitation light streams and separate them from the emission light. The MiniCube was connected to a fiber optic rotary joint (Doric Lenses) connected to the cannula. A RZ5P lock-in digital processor controlled by the Synapse software (Tucker-Davis Technologies, TDT, USA), commanded the voltage signal sent to the emitting LEDs via the LED driver (Doric Lenses). The light power before entering the implanted cannula was measured with a power meter (PM100USB, Thorlabs) before the beginning of each recording session. The light intensity to capture fluorescence emitted by 465 nm excitation was between 25-40 µW, for the 405 nm excitation this was between 10-20 µW at the tip of the fiber. The fluorescence emitted by the GCaMP6f activation in response to light excitation was collected by a femtowatt photoreceiver module (Doric Lenses) through the same fiber patch cord. The signal was then received by the RZ5P processor (TDT). On-line real time demodulation of the fluorescence due to the 405 nm and 465 nm excitations was performed by the Synapse software (TDT). A camera was synchronized with the recording using the Synapse software. Signals were exported to MATLAB R2016b (Mathworks) and analyzed offline. After careful visual examination of all trials, they were clean of artifacts in these time intervals. The timing of events was extracted from the video. For each session, signal analysis was performed on two-time intervals: one extending from –4 to 0 sec (before entering the reinforced arm) and the other from 0 to +4 sec (reinforced arm). From a reference window (from −180 to −60 sec), a least-squares linear fit was applied to the 405 nm signal to align it to the 465 nm signal, producing a fitted 405 nm signal. This was then used to calculate the ΔF/F that was used to normalize the 465 nm signal during the test window as follows: ΔF/F = (465 nm signaltest - fitted 405 nm signalref)/fitted 405 nm signalref. To compare signal variations between the two conditions (before vs after entering the reinforced arm), for each mouse, the value corresponding to the entry point of the animal in the reinforced arm was set at zero.

### Indirect calorimetry analysis

All mice were monitored for metabolic efficiency (Labmaster, TSE Systems GmbH, Bad Homburg, Germany). After an initial period of acclimation in the calorimetry cages of at least two days, food and water intake, whole energy expenditure (EE), oxygen consumption and carbon dioxide production, respiratory quotient (RQ=VCO2/VO2, where V is volume) and locomotor activity were recorded as previously described83. Additionally, fatty acid oxidation was calculated as previously reported83. Reported data are the results of the average of the last three days of recording. Before and after indirect calorimetry assessment, body mass composition was analyzed using an Echo Medical systems’ EchoMRI (Whole Body Composition Analyzers, EchoMRI, Houston, USA).

### Ex-vivo calcium imaging

Male Aldh1-L1-Cre+/− or C57BL/6J mice previously injected with GCamP6f and DREADDs viral constructs, and GCaMP6f/Glast-CreERT2 mice were terminally anaesthetized using isoflurane. Brains were removed and placed in ice-cold oxygenated slicing artificial cerebrospinal solution (aCSF, 30mM NaCl, 4.5mM KCl, 1.2mM NaH2PO4, 1mM MgCl2, 26mM NaHCO3, and 10mM D-Glucose and 194mM Sucrose) and subsequently cut into 300-µm thick PVN coronal slices using a vibratome (Leica VT1200S, Nussloch, Germany). Next, brain slices were recovered in aCSF (124mM NaCl, 4.5mM KCl, 1.2mM NaH2PO4, 1mM MgCl2, 2mM CaCl2, 26mM NaHCO3, and 10mM D-Glucose) at 37 °C for 60 minutes. Imaging was carried out at room temperature under constant perfusion (∼3 ml/min) of oxygenated aCSF. The overall cellular fluorescence of astrocytes expressing GCaMP6f was collected by epifluorescence illumination. A narrow-band monochromator light source (Polychrome II, TILL Photonics, Germany) was directly coupled to the imaging objective via an optical fiber. Fluorescence signal was collected with a 40x 0.8NA or a 63x 1.0NA water immersion objective (Zeiss, Germany) and a digital electron-multiplying charge-coupled device (EMCCD Cascade 512B, Photometrics, Birmingham, UK) as previously described (Pham et al., 2020)(Pham, 2020). A double-band dichroic/filter set was used to reflect the excitation wavelength (470 nm) to slices and filter the emitted GCaMP6 green fluorescence (Di03-R488/561-t3; FF01-523/610, Semrock). The same filter was used for slices expressing both GCaMP6f and DREADD-mCherry. Striatal slices were transferred to the imaging chamber, where 3-minute astrocyte spontaneous activity recordings were performed in slices of GCaMP6f/Glast-CreERT2 mice. In the case of striatal slices of Aldh1-L1-Cre+/− and C57BL/6J mice, we performed a basal epifluorescence recording (60 seconds), followed by a 120 second bath application of CNO (10µM) or Glutmate (30µM) and 240 seconds recording over the washing of the compounds.

The responsive regions displaying Ca^2+^ signals were scrutinized by the three-dimensional spatio-temporal correlation screening method (Pham et al., 2020). Background signal was subtracted from the raw images by using the minimal intensity projection of the entire stack. Ca^2+^ signals of individual responsive regions were normalized as dF/F0, with F0 representing the baseline intensity and quantified using Matlab (The MathWorks, France) and Igor Pro (Wavemetrics, USA). We gauged signal strength of Ca^2+^ traces of single responsive regions by calculating their temporal integration and normalizing per minute. The global temporal synchronization of detected Ca^2+^ signals was determined by the temporal Pearson’s correlation coefficients of all combinations between single Ca^2+^ regions (Pham et al., 2020).

### Brain tissue Immunofluorescence

Mice were euthanized with pentobarbital (500 mg/kg, Dolethal, Vetoquinol, France) and transcardially perfused with 0.1 M sodium phosphate buffer (PBS, pH 7.5) followed by 4% paraformaldehyde in phosphate buffer (0.1 M, pH 7.2). Brains were removed and post-fixed overnight in 4% paraformaldehyde. Afterwards, the brains were transferred to 30% sucrose in PBS for 2 days for cryoprotection. Next, 30 µm brain sections were cut in a freezing cryostat (Leica, Wetzlar, Germany) and further processed for immunofluorescence following the procedure previously described (Berland et al., 2020). Free-floating brain sections were incubated at 4°C overnight with mouse anti-Glial fibrillary acidic protein (GFAP, 1:1000, Sigma-Aldrich, Saint-Louis, USA) or mCherry (ab125096; 1:1000, Abcam, Cambridge, MA) primary antibodies. The next day, sections were rinsed in Tris-buffered saline (TBS, 0.25M Tris and 0.5M NaCl, pH 7.5) and incubated for 2 hours with secondary antibodies (1:1000, Thermo fisher Scientific, MA, USA) conjugated with fluorescent dyes: goat anti-chicken Alexa 488, donkey anti-rabbit Alexa 594, donkey anti-mouse Alexa 488 and donkey anti-rabbit Alexa 647. After rinsing, the sections were mounted and coverslipped with DAPI (Vectashield, Burlingade, California, USA) and examined with a confocal laser scanning microscope (Zeiss LSM 510, Oberkochen, Germany) with a color digital camera and AxioVision 3.0 imaging software.

### Statistical analyses

Compiled data are always reported and represented as mean ± s.e.m., with single data points plotted. Data were statistically analyzed with GraphPad Prism 9. Normal distribution was tested with Shapiro-Wilk test. When n was > 7 and normality test passed, data were analyzed with Student’s t test, one-way ANOVA, two-way ANOVA or repeated-measures ANOVA, as applicable and Holm-Sidak’s post-hoc tests for two by two comparisons. Otherwise non-parametric Mann-Whitney test. All tests were two-tailed. Significance was considered as p < 0.05.

