## Supplemental information for "Dual role of striatal astrocytes in behavioral flexibility and metabolism in the context of obesity"

**Supplementary figures legends**

**Montalban *et. al***

**List of supplementary figures**

**Supplementary figure 1.** Astrocyte Gq DREADD activation by CNO in the striatum enhances astrocyte Ca2+ activity and impact the effect of D1R agonist and D2R antagonist.

**Supplementary figure 2.** Glutamate enhances the activity synchrony and intensity of striatal neurons.

**Supplementary figure 3.** Astrocyte DREADD activation in the DS enhances neuronal Ca^2+^ signal synchrony and intensity in obese Aldh1l1-cre mice

**Supplementary figure 4**. Obesity impairs reversal learning in a procedural learning paradigm.

**Supplementary figure 5**. Consequences of astrocytes activation in the NAc on the effect of D1R agonist and D2R antagonist

**Supplementary figure 6**. Consequences of astrocytes activation in the DS on energy metabolism

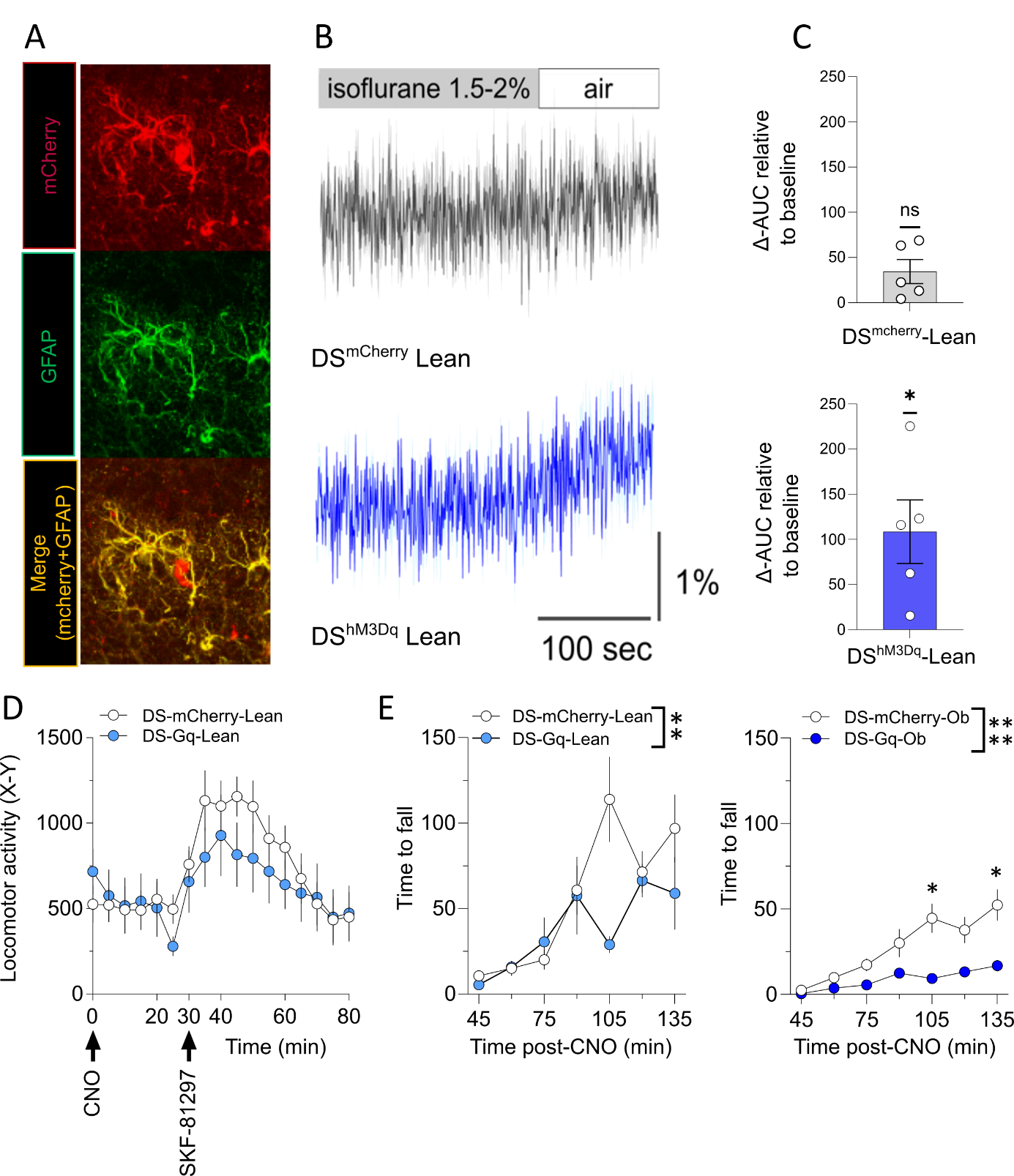

**Supplementary figure 1. Consequences of astrocytes activation in the DS**

**A** Controls of transduction of astrocytes with an AAV5 vector containing the reporter mCherry in Aldh1L1-cre mice. Coronal brain slice showing the injection site and colocalization of mCherry (red) and GFAP (green) immunostaining. **B-C** Fiber photometry Ca^2+^ signal recorded in astrocytes in vivo in Aldh1L1-cre mice co-injected with Cre-dependant Gq-DREADD and GCaMP6f viruses. Recordings were performed during the wakening of mice slightly anesthetized with isoflurane (1.5-2%). **B** Average traces of the Ca^2+^ signal under isoflurane and after isoflurane ends (n=5 for each group). **C** Difference between the AUC calculated during the 100 seconds period following the end of isoflurane and a 100 seconds period during isoflurane (from -200 to -100 sec before the end of isoflurane). Ca^2+^ increase was significant in DS^hM3Dq^ mice (One-sample t-test *p=0.0367) but not in DS^mCherry^ control mice (One-sample t-test p=0.0601; ns). **D**-Effect of astrocytes activation in the DS (CNO (0.6 mg/Kg)) on SKF-87297 induced locomotor activity (beam breaks, X-Y) in lean mice. **E** The effects astrocytes activation on DRD2 function were investigated by evaluating the immobility 45-150 min after haloperidol injection (0.1 mg.kg^-1^, i.p.), in mice treated with CNO 15 min after haloperidol injection (0.6 mg.kg^-1^, i.p.) Two-way ANOVA, Interaction F (6, 78) = 3,275 ** p=0.0063, n=8-7. **F** Same as **E** in obese DS^mCherry^ and DS^hM3Dq^ mice, Two-Way ANOVA F (2.973, 44.09) = 15.61, Sidaks post-hoc test *<0.05 n=8-9.

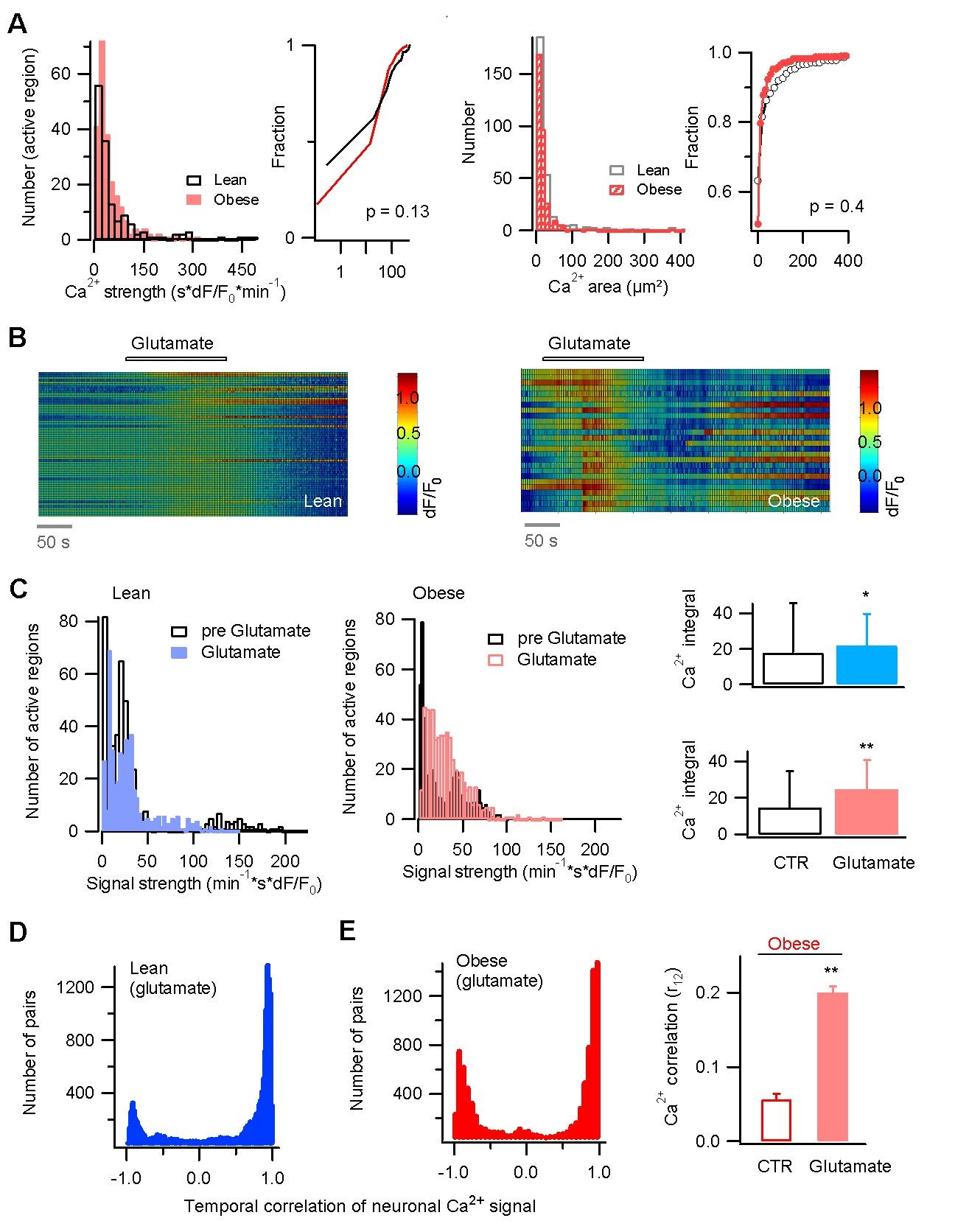

**Supplementary figure 2. Glutamate enhances the activity synchrony and intensity of striatal neurons.**

**A** Comparison of neuronal Ca^2+^ signals between lean and obese mice (Lean, 156 - 295 regions, 8 slices, 4 mice; Obese: 236 - 334 regions, 5 slices, 3 mice; Ca^2+^ strength, p = 0.4, h = 0, stats = [zval: -0.7709, ranksum: 29807]; Ca^2+^ area, p = 0.45, h = 0, stats = [zval: -0.7622, ranksum: 9.12 × 10^4^]). **B** Neuronal activity synchronously augmented by glutamate application, reflected by GCaMP6 fluorescence increase in lean and obese mice. **C** Glutamate enhances neuronal activity in lean (p = 1.8855 × 10^-4^, h = 1, stats = [zval: -3.7339, ranksum: 189018], 450 active regions, 8 slices from 4 mice) and obese mice (p = 1.6376 × 10^-8^, h = 1, stats = [zval: -5.6465, ranksum: 2.7918e+005]; 556 regions, 7 slices from 3 mice).

**D-E** Glutamate augments the temporal correlation of Ca^2+^ signals in striatal neurons. Compared to control obese mice (Fig 3B), glutamate application enhances in obese mice the temporal correlation of single Ca^2+^ signals (p = 1.12 × 10^-41^, h = 1, stats = [zval: 12.8037, ranksum: 97961689]). Obese, n = 8927 Ca^2+^ signal pairs, 3 mice; lean, n = 14415 signal pairs, 4 mice.

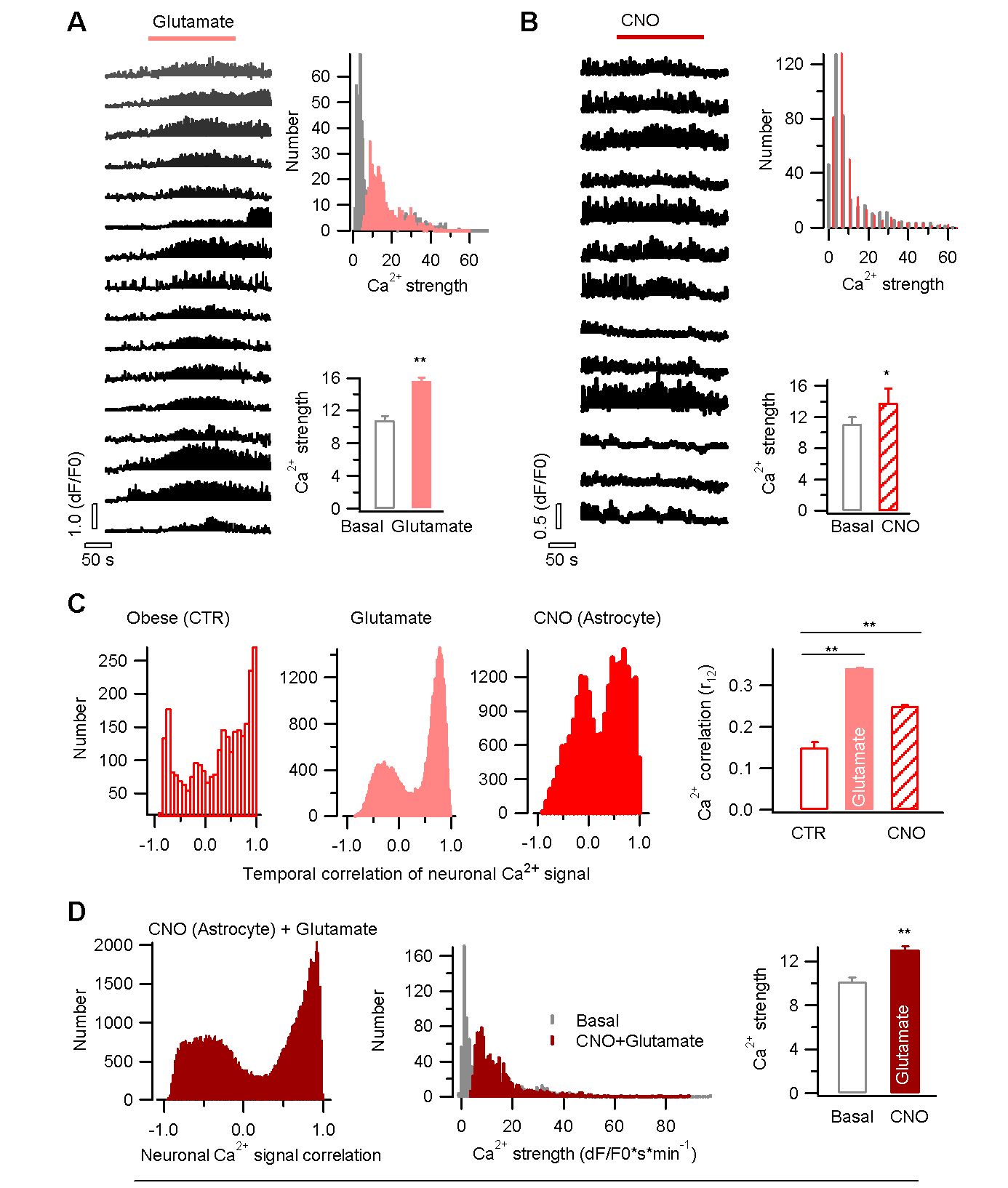

**Supplementary figure 3. Astrocyte DREADD activation in the DS enhances neuronal Ca^2+^ signal synchrony and intensity in obese Aldh1l1-cre mice.**

**A** Glutamate enhanced Ca^2+^ intensity in Aldh1l1-cre mice where hM3Dq DREADD is expressed in astrocytes, and GCaMP6 in neurons by co-injecting the vector AAV-synapsin-GCaMP6. A significant increase was observed upon glutamate application (p = 4.56 × 10^-39^, h = 1, stats = [-13.0753, 236637]; 553 signal regions, 6 slices, 4 mice; comparison made between Ca^2+^ signals before and after glutamate application). **B** Activation by CNO of astrocytes enhanced neuronal Ca^2+^ signal intensity (p = 0.043, h = 1, stats = [-2.0177, 126953]; 364 signal regions, 5 slices, 4 mice; comparison made between Ca^2+^ signals before and after CNO application). **C** Activation by CNO of astrocytes enhanced overall temporal correlation of neuronal Ca^2+^ signal in obese mice (CTR, 2492 signal pairs, three mice; glutamate, 42672 signal pairs from four mice, Wilcoxon rank sum (Mann-Whitney) test, p = 8.495 × 10^-33^ , h = 1, stats = [zval: -11.9276, ranksum: 4.873 × 10^7^]; CNO, 21952 signal pairs from four mice, Wilcoxon rank sum (Mann-Whitney) test, p = 2.97 × 10^-4^ , h = 1, stats = [zval: -3.6183, ranksum: 29250612]). **D** CNO and glutamate combined shifted the temporal correlation of Ca^2+^ signals to the positive side (73462 signal pairs from 6 mice) and augmented their strengths in striatal neurons (952 signal regions from 6 mice; comparison made between Ca^2+^ signals before and during CNO + glutamate application, Wilcoxon rank sum (Mann-Whitney) test, p = 3.7 × 10^-24^, h = 1, stats = [zval: -10.139, ranksum: 776623]).

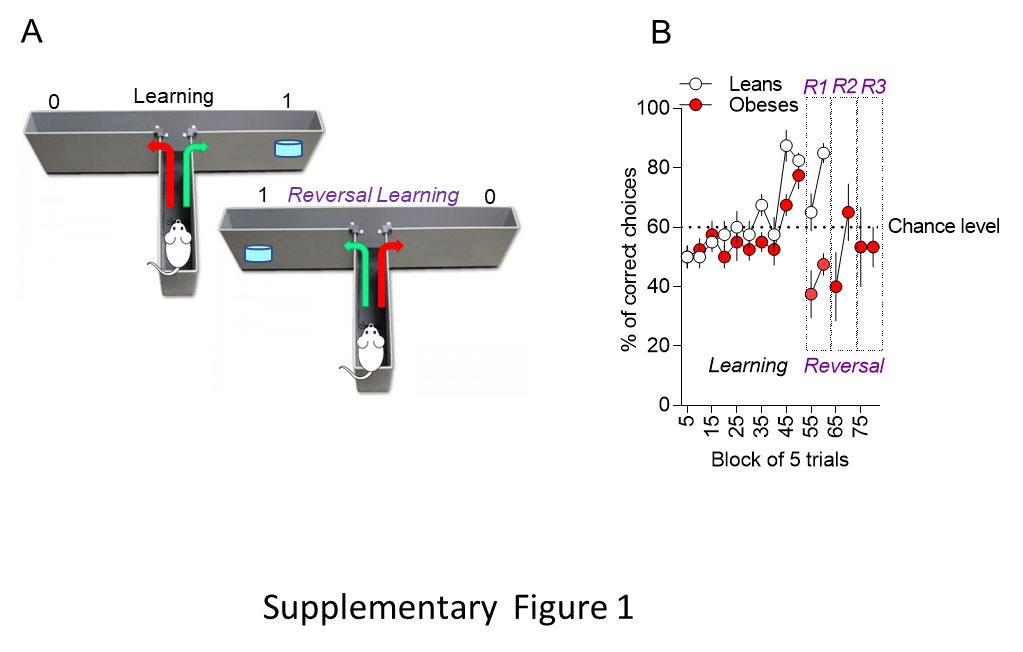

**Supplementary figure 4. Obesity impairs reversal learning in a procedural learning paradigm.**

**A**. Schematic representation of the paradigm used for learning and reversal learning. Briefly, mice were trained to find the bated arm of a maze by using an egocentric strategy. The learning phase was followed by a reversal learning, in which the bated arm was inversed. **B**. While lean mice were able to relearn the location of the bated arm already after 10 sessions (white dots), reversal was completely impaired in DIO even after 3 consecutive days of learning (6 sessions).

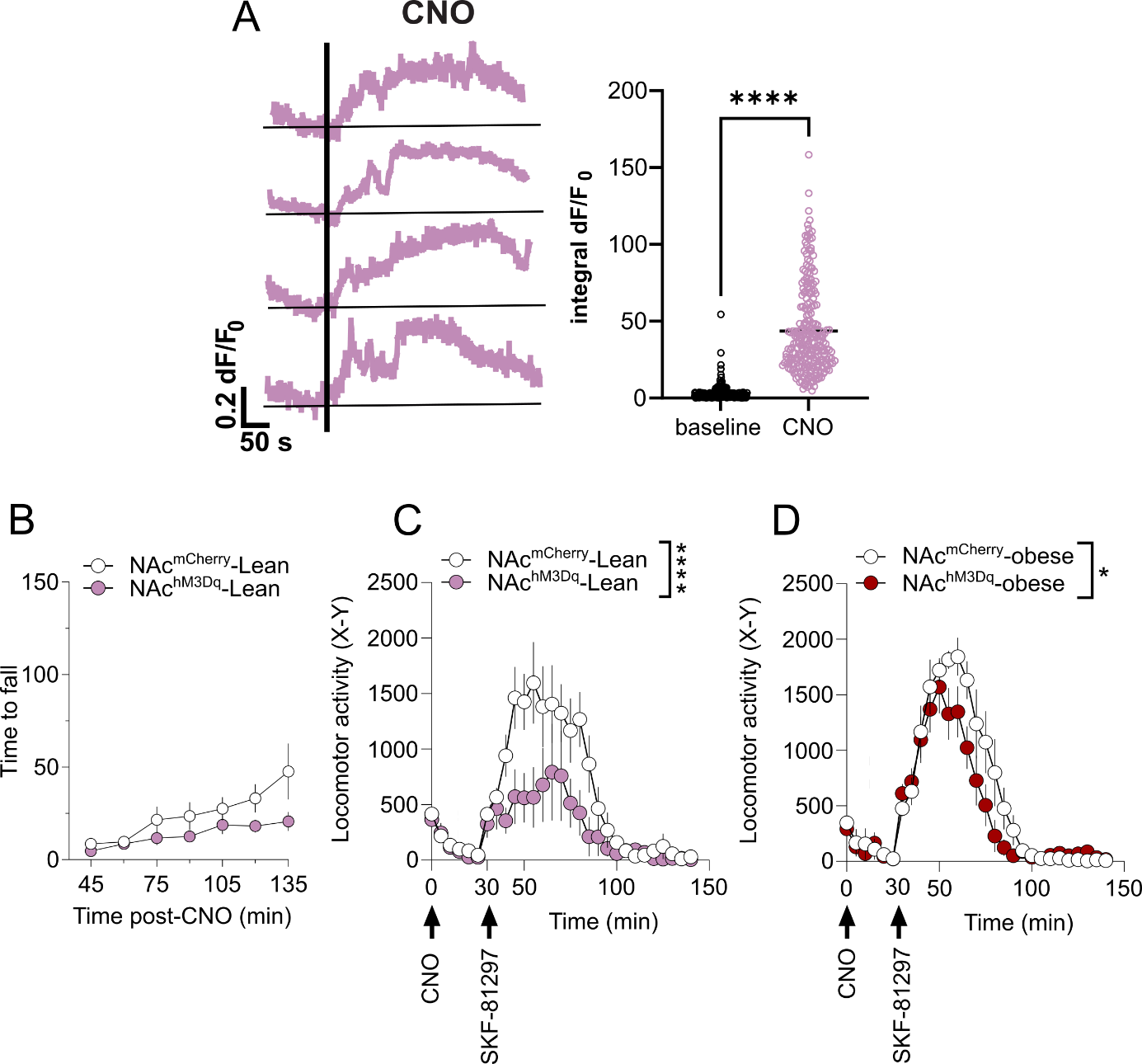

**Supplementary figure 5. Consequences of astrocytes activation in the NAc**

**A** Time-course of Ca^2+^ signal intensity increase following CNO application and integral ∆F/F_0_ during confocal imaging in slice of Aldh1L1-cre mice co-injected with Cre-dependant Gq-DREADD and GCaMP6f viruses (3 mice, 12 slices, unpaired t-test p<0.0001). **B** The consequences of astrocytes activation on NAc function were investigated in lean mice by evaluating the immobility 45-150 min after haloperidol injection (0.1 mg.kg^-1^, i.p.), in mice treated with CNO 15 min after haloperidol injection (0.6 mg.kg^-1^, i.p.) Two-way ANOVA, Interaction F (6, 60) = 3,275 p=0.1563 (ns), n=8-7. **C** Effect of astrocytes activation in the NAc (CNO (0.6 mg/Kg)) on SKF-81297 (3mg/kg) induced locomotor activity (beam breaks, X-Y) in lean mice. Two-way ANOVA, Interaction F(28, 280)=2.588, p<0.0001 n=6/group **D** Same as **C** in obese NAc^mCherry^ and NAc^hM3Dq^ mice. Two-way ANOVA, Interaction F(28, 280)=1.724 p=0.052 n=6/group.

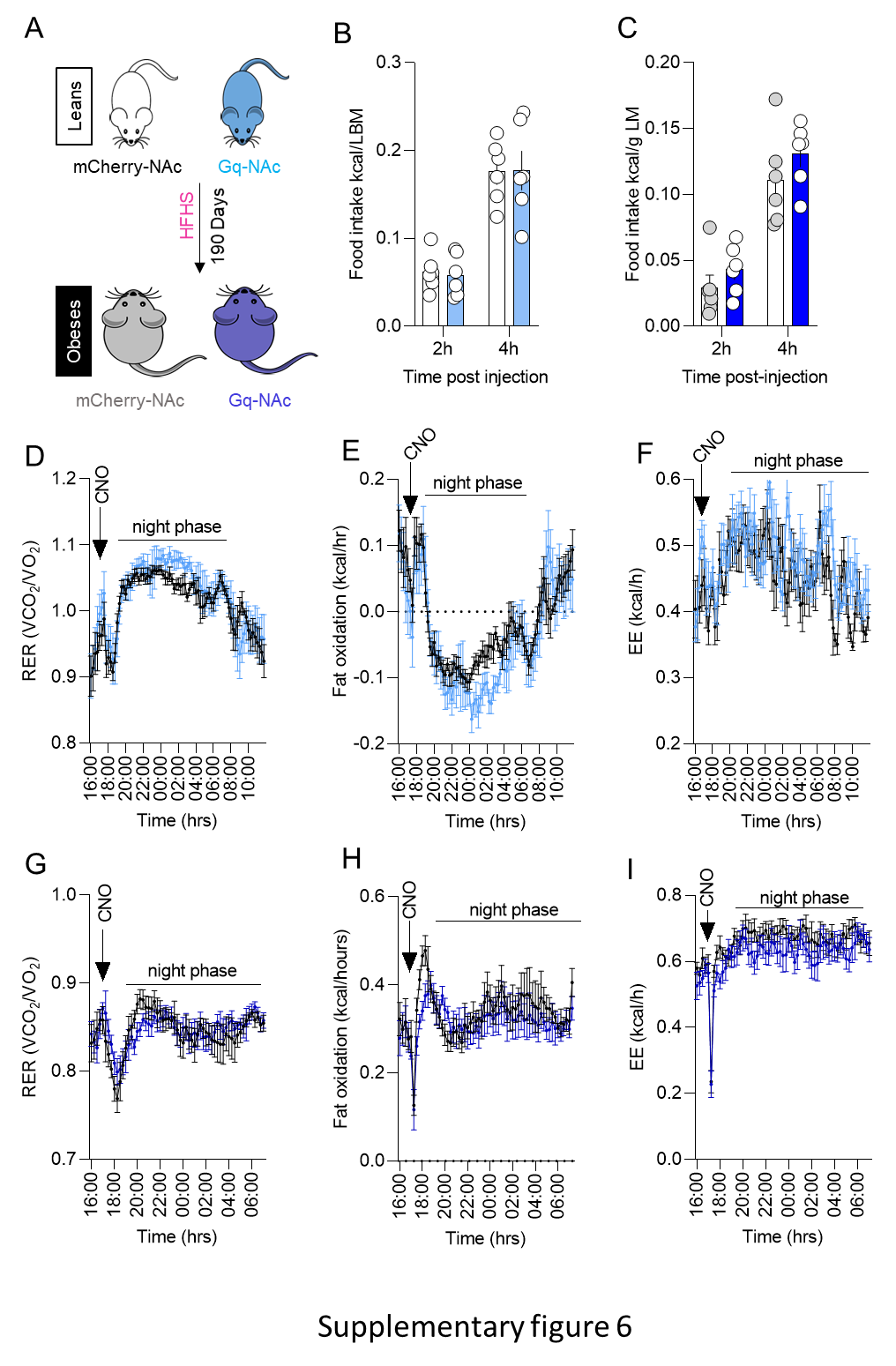

**Supplementary figure 6.** **Consequences of astrocytes activation in the DS on energy metabolism**

**A**- Schematic representation of the paradigm used for obese mice. **B**. activation of astrocytes in the DS of lean mice did not result in any changes in caloric intake. **C** Same as B for obese mice. Astrocytes activation in the DS of lean mice did not alter RER **D**, fatty acid oxidation **E** or energy expenditure **F**. **G-H-I**, same as in D-E-F for obese mice.
